# Platelets fuel the inflammasome activation of innate immune cells

**DOI:** 10.1101/800771

**Authors:** Verena Rolfes, Lucas S. Ribeiro, Ibrahim Hawwari, Lisa Böttcher, Nathalia Rosero, Salie Maasewerd, Marina Lima S. Santos, Susanne V. Schmidt, Maximilian Rothe, H. James Stunden, Luzia H. Carvalho, Cor J. Fontes, Moshe Arditi, Eicke Latz, Bernardo S. Franklin

## Abstract

The inflammasomes control the bioactivity of pro-inflammatory cytokines of the interleukin (IL)-1 family. The inflammasome assembled by NLRP3 has been predominantly studied in homogenous cell populations in vitro, neglecting the influence of cellular interactions that occur in vivo. Here, we show that platelets, the second most abundant cells in the blood, boost the inflammasome capacity of human macrophages and neutrophils, and are critical for IL-1 production by monocytes. Platelets license NLRP3 transcription, thereby enhancing ASC nucleation, caspase-1 activity, and IL-1β maturation. Platelet depletion attenuated LPS-induced IL-1β in vivo, and platelet counts correlate with plasma concentrations of IL-1β in malaria patients. Furthermore, a platelet gene signature was enriched among the highest expressed transcripts in IL-1β-driven autoinflammatory diseases. The platelet-mediated enhancement of inflammasome activation was independent of cell-to-cell contacts, platelet-derived lipid mediators, purines, nucleic acids and a host of platelet cytokines, and involved the triggering of calcium sensing receptors on macrophages by a calcium-dependent protein commonly released by platelets and megakaryocytes. Finally, we report that platelets provide an additional layer of regulation of inflammasomes in vivo.

## INTRODUCTION

An unbalanced production of the pro-inflammatory cytokines of the Interleukin-1 (IL-1) family underlies the immunopathology of several auto-inflammatory diseases. As nearly all cells express the IL-1 receptor (IL-1R), IL-1 cytokines have the ability to influence both innate and adaptive immune responses and exert broad effects in the body (Dinarello, 2009). IL-1β is unique in the medical literature: while nearly all human inflammatory diseases are caused by a host of cooperative pro-inflammatory factors, mutations in genes controlling the expression of IL-1β cause a spectrum of life-threatening auto-inflammatory syndromes (Broderick et al., 2015). Monotherapies blocking IL-1β activity in patients with auto-inflammatory syndromes result in a rapid and sustained reversal of symptoms and severity (Dinarello et al., 2012), and are currently the first line of intervention against these conditions. A growing bulk of evidence has now added other common inflammatory and metabolic conditions into the list of diseases that are responsive to IL-1β neutralization (Dinarello, 2018; Dinarello and van der Meer, 2013). This was further validated by the recent results of the Canakinumab Anti-Inflammatory Thrombosis Outcome Study (CANTOS), which showed that Canakinumab, a humanized anti-IL-1β monoclonal antibody, significantly reduced the risk for recurrent cardiovascular events (Ridker et al., 2017a), and suggested that IL-1β is associated with increased incidence of fatal lung cancer (Ridker et al., 2017b).

The expression of IL-1 cytokines is tightly regulated. For instance, the production of some members of this family (IL-1β and IL-18) is restricted to immune cells. Furthermore, upon induction, these proteins are synthesized as biologically inactive precursors in the cytosol, and a series of intracellular events are required for their maturation and release into the extracellular space. These events include the assembly of inflammasomes, intracellular multiprotein complexes formed by a sensor, such as NLRP3, the adapter molecule apoptosis-related speck-like protein containing a CARD domain (ASC), and the cysteine protease Caspase-1 (Latz et al., 2013). Upon activation, inflammasome sensors recruit ASC, which oligomerizes to form a micron-sized structure termed an ‘ASC speck’, which operate as platforms to recruit and activate Caspase-1, which processes pro-IL-1β and pro-IL-18 into their bioactive forms (Latz et al., 2013). Caspase-1 activation also drives an inflammatory lytic cell death termed pyroptosis, mediated through Gasdermin-D-induced membrane pore formation and leakage of cytosolic content (Kayagaki et al., 2015; Liu et al., 2016).

The overwhelming majority of the studies on inflammasomes were performed in vitro in monocultures of macrophages. Although this approach has led to the discovery of molecular mechanisms controlling inflammasomes, it underestimates the influence of other cell populations on these processes and in the regulation of IL-1 cytokines *in vivo*. For instance, a cooperative role for T cells for the IL-1β production by dendritic cells (DCs) has only recently been discovered (Jain et al. bioRxiv 475517).

In the last decade, platelets have been increasingly recognized of their roles in immunity (Allen et al., 2019; Dann et al., 2018; Kral et al., 2016; Passacquale et al., 2011a). Approximately one trillion platelets (150– 450 × 10^9^/L) circulate in the blood of a healthy individual, a number that surpasses all other leukocytes in the vasculature by several folds. Platelets have been reported to produce IL-1 cytokines (Allam et al., 2017; Denis et al., 2005; Thornton et al., 2010a), and more recently to assemble an NLRP3 inflammasome (Cornelius et al., 2019; Hottz et al., 2013). They could therefore be relevant cellular sources of IL-1 cytokines or extracellular ASC specks in vivo or alter the inflammatory responses of other immune cells.

Using a series of complementary techniques, human and mouse platelets and megakaryocytes, as well as transgenic (Tzeng et al., 2016b) and *knock in* inflammasome reporter mouse models, we report here that platelets, as well as megakaryocytes, do not express the components of the canonical inflammasome (NLRP3, ASC, and Caspase-1). Nevertheless, co-culture with platelets boosted the inflammasome activation and production of IL-1α, IL-1β, and IL-18 from human macrophages and neutrophils. Furthermore, platelets were crucial for the optimal production of IL-1 cytokines by human monocytes. We found that platelets influenced the inflammasome activation of these cells *in trans* through the enhancement of NLRP3 and pro-IL-1β transcription. The effect of platelets on human macrophages did not require direct cell contact, platelet-derived nucleic acids, or purines, but it could be extinguished by heat inactivation. Using a model of antibody-induced thrombocytopenia, we found that platelet depletion attenuated IL-1β responses while enhancing the systemic production of TNFα in response to LPS in vivo. Supporting these findings, blood platelet counts correlated positively with plasma levels of IL-1β in naturally infected malaria patients. Moreover, we observed an enriched platelet signature among the highest expressed genes in a cohort of pediatric patients with mutations in the NLRP3 gene causing Muckle-Wells Syndrome (MWS) and Neonatal-onset multisystem inflammatory disease (NOMID).

In summary, we show that platelets fuel inflammasome activation in immune cells and shape IL-1 inflammation. Thus platelet-modifying therapies could have widespread implications for autoinflammatory and thrombotic diseases.

## RESULTS

### Platelets boost inflammasome-driven IL-1 production in macrophages and neutrophils

The NLRP3 inflammasome is primarily assembled in myeloid cells. Although activated platelets are known to form aggregates with immune cells and modulate their function (Allen et al., 2019; Dann et al., 2018; Kral et al., 2016; Passacquale et al., 2011b) the effect of platelets on inflammasome activation in innate immune cells remains unexplored. To address this question, we investigated the influence of platelets on the production of IL-1 cytokines elicited by the *bona fide* NLRP3 inflammasome activators Nigericin and ATP in mouse and human immune cells. Mouse bone marrow-derived macrophages (BMDMs) (**Figure 1A**), human monocyte-derived macrophages (hMDMs) (**Figure 1B**), human neutrophils (**Figure 1C**) and human CD14^+^ monocytes (**Figure 1D**) were cultured alone or in the presence of increasing concentrations of platelets. Co-cultures were then left untreated or primed with LPS, followed by NLRP3 activation with Nigericin or ATP. The levels of IL-1β and TNFα were measured in cell-free supernatants by homogeneous time-resolved fluorescence (HTRF). A multiplex cytokine assay was used to additionally measure the production of IL-1α and IL-18, two other members of the IL-1 family, along with several other cytokines and growth factors in hMDMs and human neutrophils (**Figure S1A-B**). Remarkably, co-culture with platelets boosted the production of IL-1β from both mouse and human inflammasome-activated macrophages, as well as from human neutrophils in a concentration-dependent manner (Figure 1A-C). Addition of platelets also boosted the release of IL-18 and IL-1α from activated hMDMs (**Figure S1A**). Extending these results, we found that the addition of platelets to human neutrophils enhanced their production of IL-8 (**Figure S1B**). Platelets cultured alone produced RANTES (CCL5) and low levels of IL-18, however, those were close to the detection limit of our assays. Platelets lacked expression of most of the other investigated cytokines (**Figure 1** and **S1A-B**), indicating that they do not directly contribute to the cytokines measured in the co-cultures. Notably, the addition of platelets to mouse macrophages, human monocytes, and neutrophils, but not human macrophages, resulted in diminished TNFα production in response to LPS stimulation. These results suggest that despite NFkB being a common transcription factor regulating the production of TNFα and IL-1 cytokines, platelets exert a selective effect on the production of inflammasome-governed cytokines.

**Figure 1.**
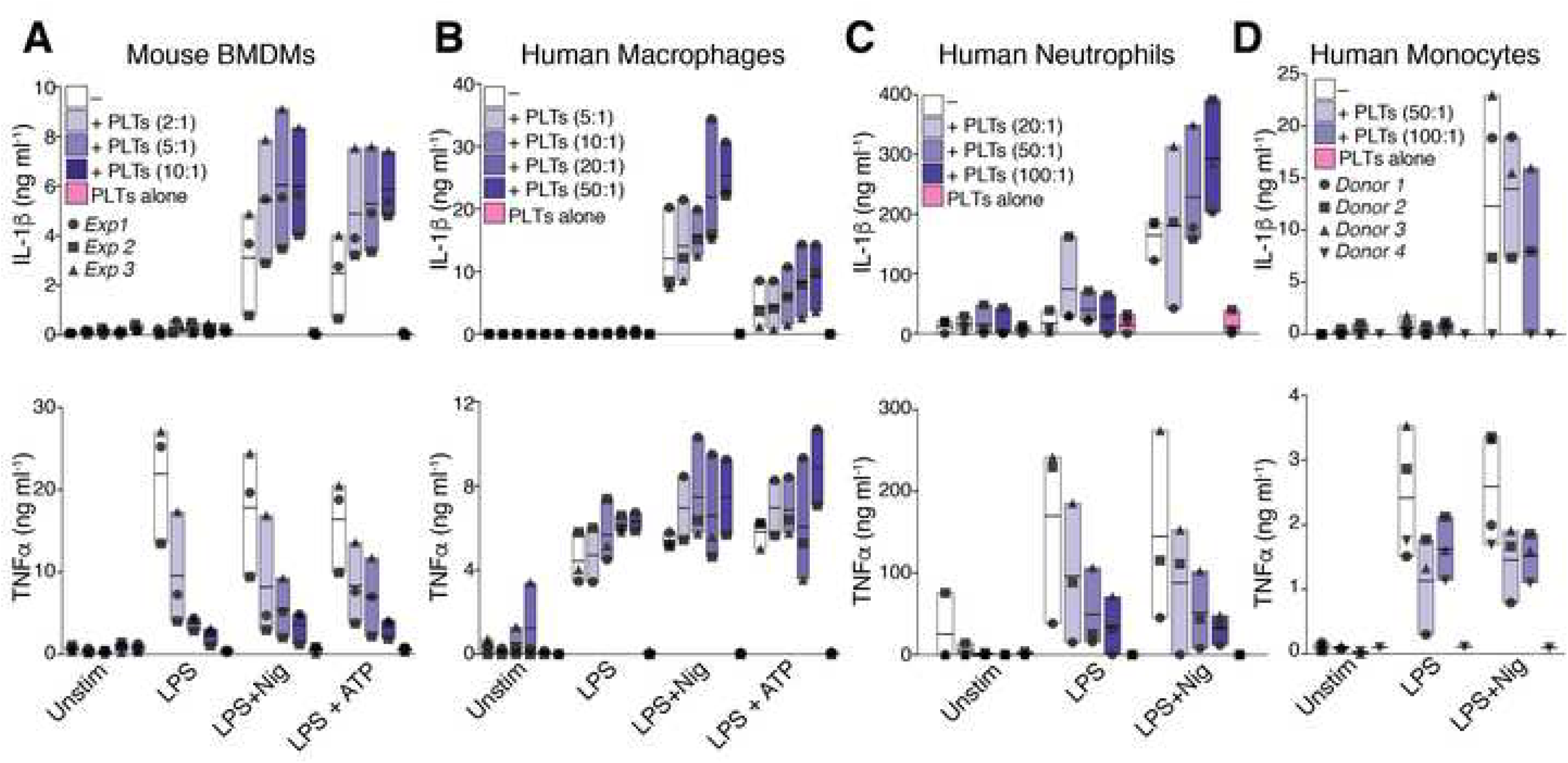
Platelets amplify inflammasome activation of innate immune cells. (**a**) IL-1β and TNFα levels in cell-free supernatants of unstimulated (Unstim), or LPS-primed (200ng/mL, 3 hours), and nigericin (10µM, 90 min), or ATP (5mM, 90 min)-activated wild-type BMDMs. Cells were cultivated alone (−) or in the presence of platelets (+PTLs) in the indicated ratios (platelets: macrophage). Each symbol represents the average of triplicate cell cultures of individual mice (n = 3 independent experiments). (**B**) IL-1β and TNFα levels in cell-free supernatants of human monocyte-derived macrophages (hMDMs, n = 3), (**C**) human neutrophils (n = 3), or (**D**) human CD14^+^ isolated monocytes (n = 4) stimulated as in **a**. Floating bars (with mean and minimum to maximum values) are shown from pooled data from independent experiments with cells and platelets from different donors. Each symbol represents the average from technical triplicates per donor, or mice. (See also **Figure S1**).

Next we asked whether platelets also influence the activity of other inflammasomes. To test this, we activated the NLRC4 inflammasome in co-cultures of human platelets and macrophages with a plasmid encoding the T3SS apparatus (rod protein) from *Salmonella typhimurium* (PrgI), together with a protective antigen protein (PA) (Zhao et al., 2011). Stimulation of the NLRC4 inflammasome in human macrophages induced robust IL-1β levels which were further enhanced by the addition of platelets (**Figure S1C**), indicating that the effect of platelets was not specific to the NLRP3 inflammasome. Co-culture with platelets also boosted IL-1β production from human macrophages that were primed with TLR2 or TLR7/8 agonists (Pam3cysk4 and R848, respectively) (**Figure S1D**), indicating that the platelet effect is not exclusively mediated through TLR4. However, blockade of TLR4 signaling on hMDMs with Resatorvid (TAK242), a small-molecule inhibitor of TLR4 (Matsunaga et al., 2011), partially prevented the effect of platelets (**Figure S1E**), indicating that the platelet effect on hMDMs is in part orchestrated by TLR4. Together these data show that platelets boost the production of IL-1 cytokines in inflammasome-activated innate immune cells.

### Platelets are critical for the optimal production of IL-1 cytokines by human monocytes

Notably, the addition of platelets did not influence the production of IL-1β by inflammasome-activated human monocytes (**Figure 1D**). To determine whether this was due to the steady-state presence of contaminating platelets, we performed platelet depletion experiments. CD14^+^ monocytes were isolated from fresh peripheral blood using commercially available monocyte isolation kits, with or without the addition of a platelet removal antibody cocktail, and purity was assessed by flow cytometry (**Figure 2A**). The addition of the platelet removal component efficiently reduced the number of contaminating free platelets (CD41^+^, CD14^−^), as well as the frequency of platelet-monocyte aggregates (CD41^+^, CD14^+^), while enriching the frequency of platelet-free monocytes (CD14^+^, CD41^−^) (**Figure 2A and 2B**). Importantly, the platelet removal component mainly comprises antibodies in PBS, which did not affect lactate dehydrogenase (LDH) release, a marker for cellular toxicity and cytolysis (**Figure S2A**). Removal of platelets from isolated CD14^+^ monocytes impaired their ability to release most IL-1 cytokines upon inflammasome activation, and this could be ameliorated by re-addition of freshly isolated autologous platelets (**Figure 2C** and **Figure S2B**). These data indicate that platelets are crucial for monocytes to trigger a maximal inflammasome response.

**Figure 2.**
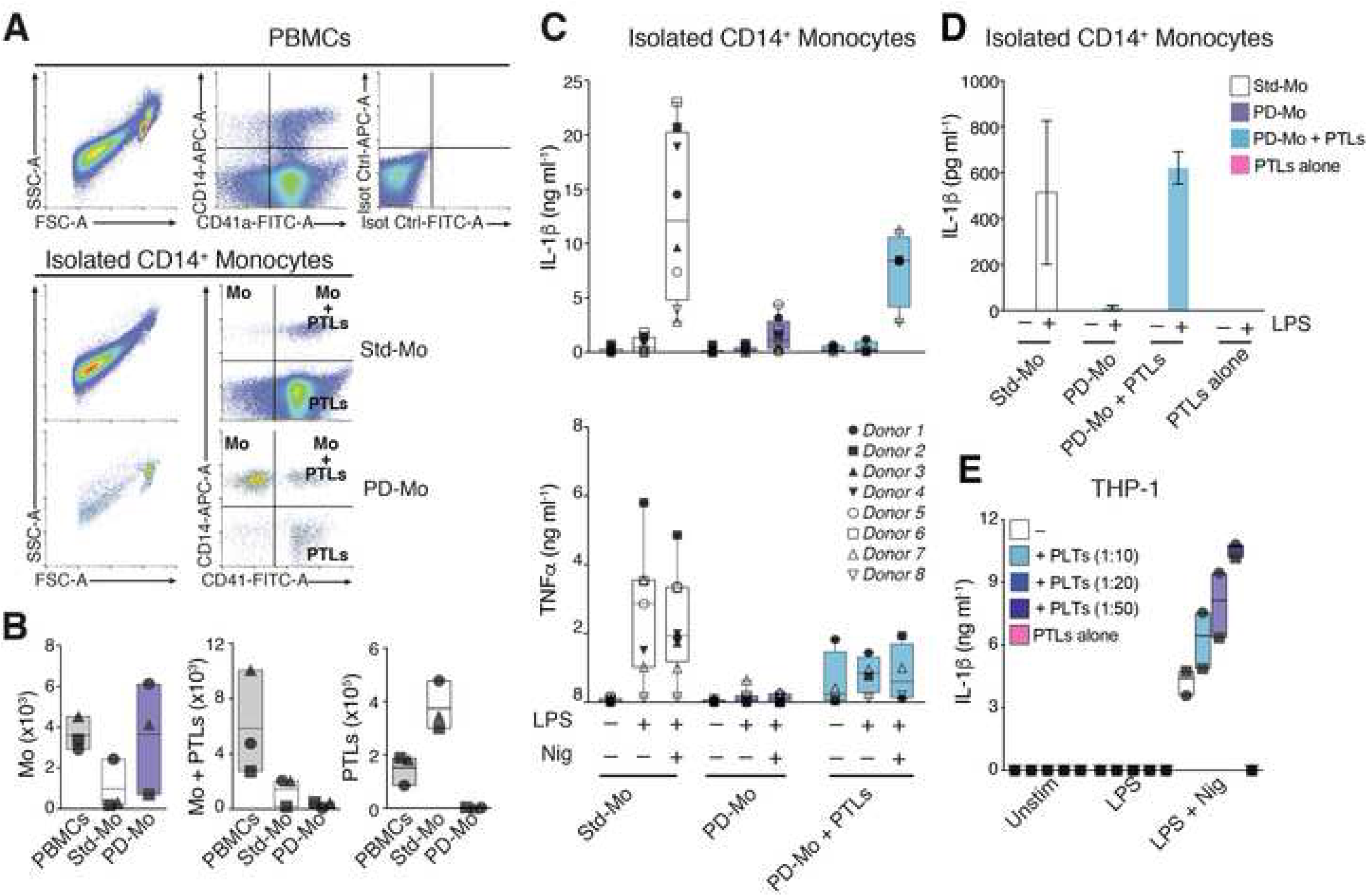
Platelets are critical for the production of IL-1 cytokines by human primary monocytes. (**a**) Representative flow cytometry scatter characteristics of human PBMCs, or CD14^+^ monocytes that were isolated with standard magnetic separation kits (Standard Monocytes, Std-Mo), in the presence or absence of a commercially available platelet depletion cocktail during isolation (Platelet-depleted Monocytes, PTL-deplt-Mo). Expression of CD41 (platelet marker) and CD14, as well as staining with Isotype IgG controls was assessed in all cell populations. (**B**) Flow cytometric-based quantification of PBMCs and isolated CD14^+^ monocytes, showing the frequency of contaminating platelets (CD41^+^, CD14^+^), platelet-monocyte aggregates (CD41^+^, CD14^+^), and monocytes (CD41^−^, CD14^+^) before vs. after the removal of platelets. (**C**) IL-1β, IL-1α and IL-18 levels in cell-free supernatants of Std-Mo vs PTL-deplt-Mo, or from PTL-depth-Mo that were co-cultured with freshly isolated autologous platelets in a 50:1 ratio (platelet: monocyte, PTL-deplt-Mo + PLTs 50:1) and stimulated, as indicated, with LPS and Nigericin. (**D**) IL-1β levels in cell-free supernatants of non-canonically activated standard, platelet-depleted, or PTL-deplt monocytes that were added with freshly isolated autologous platelets. Cells were left unstimulated (Unstim), or primed with LPS (1µgml^-1^, for 16 hours). Graph shows mean ± SD from technical triplicates from one experiment. (**E**) IL-1β levels in cell-free supernatants of inflammasome-activated THP-1s cultured alone or in the presence of growing concentrations of human platelets. Graphs (**A-C**, and **E**) show floating bars (with mean and minimum to maximum values) from pooled data from several independent experiments. Each symbol represents the average of technical triplicates from different donors. See also **Figure S2**.

Human monocytes have recently been described to activate an alternative inflammasome in which LPS alone is sufficient to trigger caspase-1-dependent IL-1β maturation and secretion (Gaidt et al., 2016). To test the effect of platelet removal in alternatively-activated human primary monocytes, we stimulated standard or platelet-depleted CD14^+^ monocytes with LPS (1μg ml^-1^) for 16 hours. Similar to the effect on the canonical inflammasome (**Figure 2C**), platelet removal also extinguished the IL-1β response from alternatively activated cells (**Figure 2D**). Furthermore, re-addition of autologous platelets rescued IL-1β production in alternatively-activated monocytes (**Figure 2D**).

We next asked whether co-culture with platelets could also enhance inflammasome activation in the surrogate monocytic cell line THP-1, which have been used extensively to characterize the biology of the inflammasome. As expected (Gaidt et al., 2016, 2017), inflammasome-activated THP-1s were able to produce IL-1β irrespective of the presence of platelets. Nevertheless, co-culture boosted the production of IL-1β by THP-1 cells to a level comparable to that produced by primary human monocytes (from∼3 to ∼10 ng ml-1) (**Figure 2E**). No IL-1β was detected on platelets cultivated alone. Taken together, these findings reveal that platelets are critical for optimal inflammasome-driven IL-1 cytokine production by human monocytes.

### The influence of platelets on IL-1 responses *in vivo* and in human disease

As monocytes and macrophages are relevant tissue sources of IL-1 cytokines, we reasoned that platelets might influence IL-1β responses in vivo. To investigate this hypothesis, we induced thrombocytopenia in C57BL/6j mice by i.v. injection of 2 µg/g of body weight of a rat anti-mouse GPIba monoclonal antibody (αCD42b). After 2 hours, treatment with αCD42b resulted in a drop in blood platelet counts (**Figure 3A**). Upon LPS challenge, platelet-depleted mice exhibited a non-significant trend toward decreased IL-1β (**Figure 3B**). Corroborating previous observations (Xiang et al., 2013) and our *in vitro* findings, thrombocytopenic LPS-challenged mice had increased serum concentrations of TNFα (*P*= 0.043) and IL-6 (*P*= 0.001) (**Figure 3B**). Sera IL-18 levels remained unaffected by platelet-depletion in this model. These findings support a role for platelets in modulating cytokine levels in vivo.

**Figure 3.**
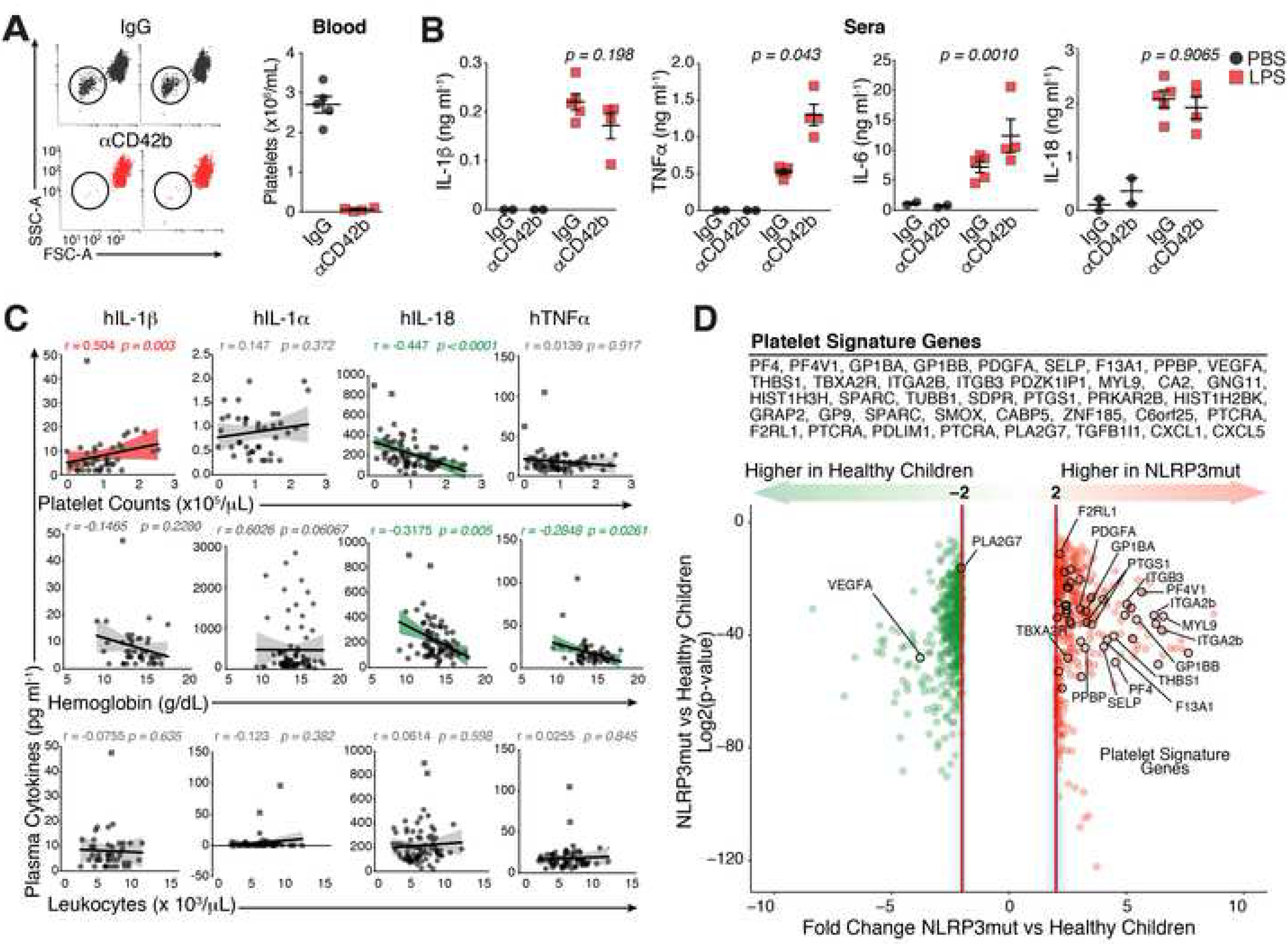
The platelet influence on cytokines levels *in vivo*. (a) Representative flow cytometry scatter plots and quantification of platelets in whole blood of C57BL/6 mice that were injected i.v. with 2 µg/g of body weight of a rat anti-mouse GPIba monoclonal antibody (αCD42b, n = 5) or control rat IgG (IgG, n = 5). (**B**) Assessment of IL-1β, IL-18, TNFα, and IL-6 levels in sera from IgG- or anti-CD42b-treated mice 2h after i.v. injection of 100 µg of LPS. Values are means ± S.E.M. Each symbol represents one mouse. (**C**) Correlation between circulating levels of IL-1α, IL-1β, IL-18 and TNFα and blood platelet, hemoglobin levels, and leukocyte counts in the plasma of naturally infected malaria patients (n= 78). Dots are semitransparent, with darker symbols indicating overlapping points. Shaded areas are based on univariate linear regressions with 95% confidence bands of the best-fit line and show positive (red) correlations. Two-tail Spearman correlation coefficient (R) and significance (*P*) are shown. Squared symbols represent patients that scored positive as outliers based on the ROUT method, and were excluded from the statistical analysis. (**D**) Volcano Plot of the expression (Log2 fold change) vs. significance (log2p value) showing the frequency of Platelet Signature Genes in a genome-wide expression profile of whole blood from healthy (n= 14) vs pediatric patients (n= 22) with mutations in the NLRP3 gene (NLRP3mut) that cause Muckle-Wells Syndrome and Neonatal-onset multisystem inflammatory disease (NOMID) from a publicly available dataset (Jr et al., 2013). The platelet gene signature was generated from direct comparisons of publicly available gene expression datasets from purified human platelets (Eicher et al., 2016; Rowley et al., 2011).

Supporting this hypothesis, a positive correlation between blood platelet counts and plasma IL-1β concentrations was recently reported in a cohort of 500 Caucasian healthy volunteers (Netea et al., 2016; Tunjungputri et al., 2018). However, blood leukocyte counts were equally correlated with plasma levels of IL-1β in that cohort, hindering the precise contribution of platelets for the plasma IL-1β concentrations described (Tunjungputri et al., 2018). To investigate the relationship platelet count and IL-1β concentrations in the context of disease, we studied a cohort of human subjects naturally infected with *Plasmodium vivax*, the predominant cause of malaria in the Brazilian Amazon basin. Infections with *P. vivax* are known to cause a strong pro-inflammatory cytokine imbalance (Andrade et al., 2010; Clark et al., 2006) with thrombocytopenia and anemia being the most commonly associated complications. We found that platelet counts were positively correlated with plasma IL-1β concentrations (Spearman**’**s R = 0.504, *P = 0.0028*). Despite a positive trend, no correlations were found between platelet count and IL-1α (R= 0.15, *P = 0.37*) or TNFα (R= 0.014, *P = 0.917*) (**Figure 3C**). Unexpectedly, plasma levels of IL-18 were negatively correlated with platelet counts (R=-0.45, *P < 0.0001*), which may suggest that different mechanisms regulate the production of this cytokine in vivo. Indeed, both IL-18 (R= −0.32, p = 0.005) and TNFα (R= −0.29, p = 0.003) correlated with hemoglobin levels (**Figure 3C**), suggesting that anemia may be a contributing factor for the regulation of these cytokines, but not for the associations between platelet counts and IL-1β. Importantly, none of the investigated cytokines correlated with leukocyte counts (**Figure 3C**) in the malaria cohort. As leukocytes are a relevant source in IL-1 cytokines in the blood, these data support that platelets are major contributors in the regulation of IL-1 cytokines levels in this context.

We next asked whether there is a relationship between platelets and inflammation in exclusively IL-1β-driven human autoinflammatory syndromes (Broderick et al., 2015). We generated a platelet gene-signature comprising 45 transcripts known to be either platelet-specific or strongly associated with platelet activity from direct comparisons of publicly available gene expression analyses of purified human platelets (Eicher et al., 2016; Rowley et al., 2011). Remarkably, 42 out of 45 platelet signature genes were upregulated in the whole blood of pediatric patients harboring mutations in NLRP3 (*NLPR3mut*) associated with the IL-1-driven autoinflammatory disorders Muckle-Wells Syndrome (MWS) and NOMID (Balow et al., 2013) compared to healthy children (**Figure 3D**). For several of these genes, more than one transcript variant was upregulated. Collectively, these findings support a role for platelets in shaping IL-1-driven inflammation in human disease.

### Platelet effect on immune cells is independent of platelet-derived IL-1 cytokines or inflammasome components

Previous reports have indicated that platelets express IL-1 cytokines, including IL-1α (Thornton et al., 2010b), IL-1β (Boilard et al., 2010; Denis et al., 2005) and IL-18 (Allam et al., 2017). However, as shown in **Figure 1** and **Figure S1A-B**, IL-1α and IL-1β were not detected in monocultures of platelets in our experimental settings. To exclude the possibility that platelets directly contribute to the IL-1 cytokines measured in co-cultures with hMDMs, we performed co-culture experiments using platelets from mice with genetic deficiency in IL-1 gene or macrophages from mice with genetic deficiency of their receptors. Firstly, we activated the NLRP3 inflammasome in wild-type BMDMs co-cultured with platelets from either wild-type or IL-1β deficient (*Il1b*^−/−^) mice. We found that *Il1b*^−/−^ platelets were equally able to boost IL-1β production in inflammasome-activated wild-type macrophages as wild-type platelets (**Figure 4A**). Similarly, *IL1r*-deficient (*Il1r*^−/−^) macrophages exhibited the expected IL-1β production in response to platelets (**Figure 4B**), thus also excluding a role for platelet-derived IL-1α in mediating the response. Finally, addition of platelets to macrophages deficient in the IL-18 receptor (*Il-18r*^−/−^) equally boosted IL-1β production in response to LPS + Nigericin (**Figure 4C**), indicating that neither platelet-derived IL-1α/β nor IL-18 are responsible for their effects of platelets on macrophage cytokine production.

**Figure 4.**
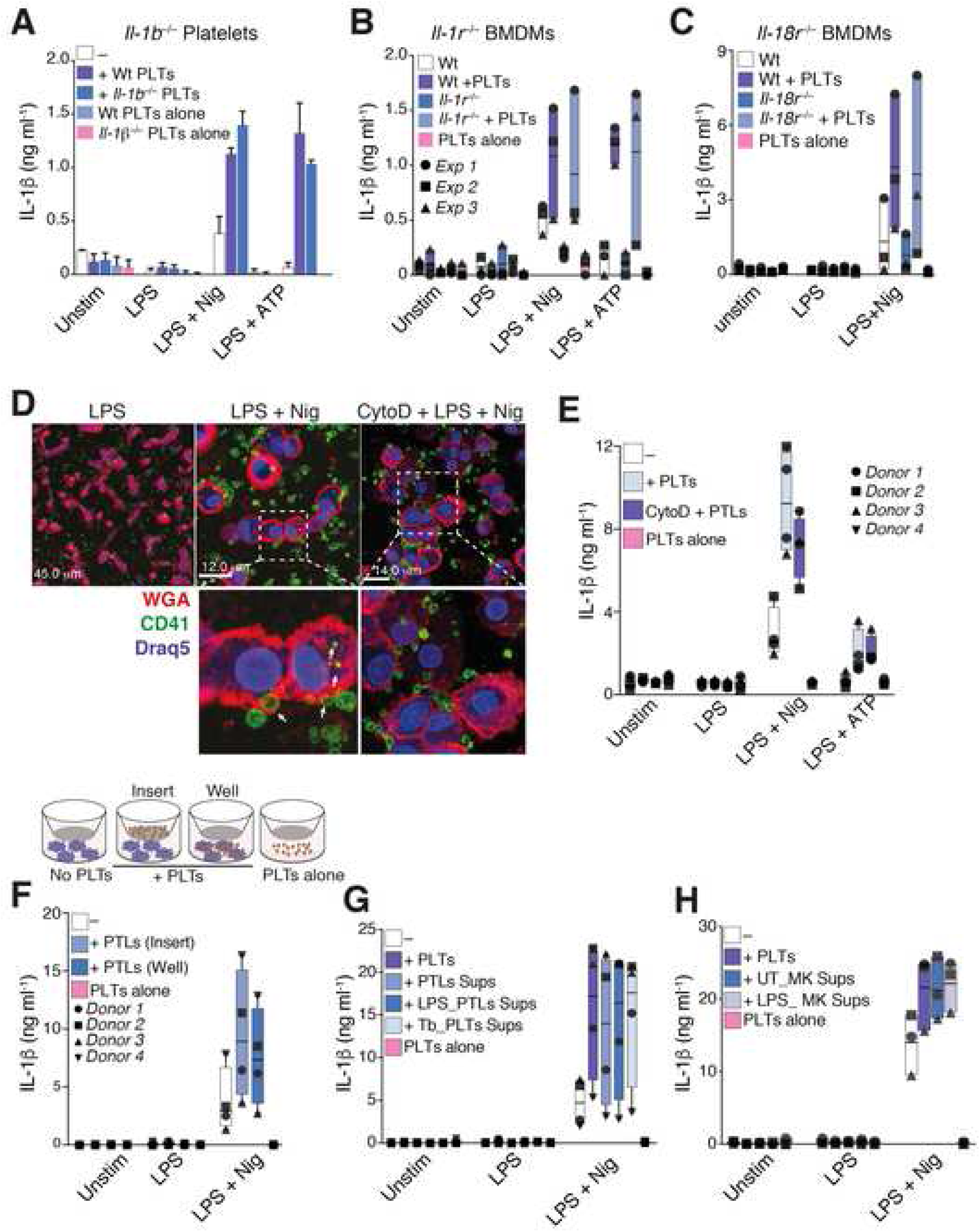
The platelet-mediated inflammasome boosting is independent of platelet-derived IL-1 cytokines, or inflammasomes. (a) HTRF measurements of mouse IL-1β in cell-free supernatants of wild-type BMDMs cultivated alone (—), or in the presence of platelets (+PTLs, 5:1 platelet-to-BMDM ratio) from wild-type, or IL-1-deficient (*Il1b*^−/−^) mice. Mean + SD from 3 technical replicates from one experiment. (**B**) HTRF measurements of mouse IL-1β in cell-free supernatants of wild-type, or IL-1R deficient (*Il1r*^−/−^), or (**C**) *Il18r*^−/−^ BMDMs cultivated alone, or in the presence of platelets (5:1 ratio) from wild-type mice. (**D**) HTRF measurements of mouse IL-1β in cell-free supernatants of wild-type, or *Nlrp3*^−/−^, or Pycard^−/−^ BMDMs cultivated alone, or in the presence of platelets (5:1 ratio) from wild-type mice. **B**-**D** Floating bars (with mean and minimum to maximum values) are shown from pooled data from three independent experiments. Each symbol represents the average of technical triplicate cultures from each mouse. (**D**) Confocal imaging of human monocyte-derived macrophages (hMDMs) that were co-cultured with platelets (50:1 ratio). Cells were pre-treated or not with Cytochalasin D (50µM, 30 min) before being added with platelets. Co-cultures were primed with LPS (200ng/mL, 3 hours) and activated with nigericin (10µM, 90 min). Blue (Draq5, nuclei), Red (WGA, plasma membrane), Green (anti-CD41, platelets). Scale bars are indicated. (**E**) HTRF measurements of IL-1β in cell-free supernatants of hMDMs treated as in d. (**F**) Schematics of a trans-well system and HTRF measurements of IL-1β levels in cell-free supernatants of co-cultures of hMDMs and platelets seeded in single cultures, direct co-cultures (Well), or trans-well cultures (Insert), separated by a 0.4 µm pore membrane. (**G**) HTRF measurements of IL-1β levels in cell-free supernatants of hMDMs cultured alone, or in the presence of platelets, or supernatants of resting, or LPS, or thrombin-activated platelets, as well as supernatants of resting, or LPS-activated megakaryocytes (**H**) Graphs show floating bars (with mean and minimum to maximum values) from pooled data from four independent experiments. Each symbol represents the average of technical triplicates from different donors.

Recent studies reported that the NLRP3 inflammasome is assembled in human platelets and that inflammasomes in platelets are involved in the pathogenesis of dengue (Hottz et al., 2014), and sickle cell disease (Vogel et al., 2018). To determine whether the simultaneous activation of platelet-NLRP3 could contribute to the production of IL-1 cytokines in co-cultures, we investigated whether addition of wild-type platelets could compensate for macrophages deficient for inflammasome components NLRP3 (*Nlrp3*^−/−^) or ASC (*Pycard*^−/−^). We stimulated LPS-primed *Nlrp3*^−/−^, or *Pycard*^−/−^ BMDMs that were cultured alone, or in the presence of platelets, with nigericin or ATP. As expected, macrophages from both *Nlrp3*^−/−^ and *Pycard*^−/−^ mice failed to produce IL-1β in response to inflammasome activation (**Figure S2C**). Addition of wild-type platelets to *Nlrp3*^−/−^ or *Pycard*^−/−^ macrophages had no effect on the IL-1β production by these cells (**Figure S2C**), indicating that activation of inflammasomes in platelets is not involved in the amplification of macrophage IL-1β responses. To determine whether the NLRP3/ASC inflammasome is expressed and assembled in platelets we imaged inflammasome assembly in total bone marrow (BM) cells from transgenic (Tg) ASC-mCitrine mice (Tzeng et al., 2016a). ASC-mCitrine^+^ BM cells were stained for leukocytes (CD45), neutrophils (Ly6G), and platelets (CD41), and assessed by flow cytometry and confocal microscopy. Whilst ASC was clearly visualized and assembled into fluorescent specks in inflammasome-activated leukocytes (CD45^+^), macrophages (CD45, F4/80^+^) and neutrophils (CD45^+^, Ly6G^+^), and extracellular inflammasomes were visible as previously described (Baroja-Mazo et al., 2014; Franklin et al., 2014a) ASC was not observed in platelets and megakaryocytes (CD41^+^cells) (**Figure S3A-B**). Similar results were observed from the imaging of total BM cells from ASC-mCherry *knock in* mice (**Figure S3C**).

We next evaluated the expression of the canonical inflammasome components in platelets and PBMCs from healthy volunteers. The purity of platelet preparations was assessed by flow cytometry, microscopy, and qPCR using platelet (PF4), or leukocyte markers (CD45and CD14) (**Figure S4A-E**). Importantly, both freshly isolated human and mouse platelets remained viable after purification and responded to thrombin stimulation by up-regulating P-selectin (CD62P) (**Figure S3D** and **S4D**). Nevertheless, purified human platelets did not express the inflammasome molecules NLRP3, ASC (PYCARD) and Caspase-1, or IL-1β either at the mRNA (**Figure S4E**) or the protein level (**Figure S4F-G**), although these molecules were promptly detected in PBMCs from the same donors, and increased upon LPS stimulation. Likewise, unlike the human monocytic cell line THP-1, no expression of IL-1β and inflammasome molecules were detected in the MEG-01 human megakaryocytic cell line (**Figure S3E**), which, compared to THP-1, failed to secrete mature IL-1β upon inflammasome activation (**Figure S3F**).

Finally, we assessed and analyzed publicly available transcription profiles of purified platelets from five independent studies (Devignot et al., 2010; Gnatenko et al., 2005; Londin et al., 2014; Raghavachari et al., 2007; Spivak et al., 2014), including the gene expression profiles of platelets from patients with Dengue (Devignot et al., 2010) and sickle cell disease (Raghavachari et al., 2007) (**Figure S5**). In all these studies we re-assessed the expression of IL-1β and the inflammasome components NLRP3, ASC and Caspase-1. The expression of platelet markers PF4 (CXCL4) or PDGFA were used as comparisons (**Figure S5**). None of the NLRP3, ASC, Caspase-1 and IL-1β transcript variants were detected in human platelets in these studies (Devignot et al., 2010; Gnatenko et al., 2005; Londin et al., 2014; Raghavachari et al., 2007; Spivak et al., 2014) (**Figure S5**). Together with our previous findings, these observations support the conclusion that both human and mouse platelets do not express the components of the canonical NLRP3 inflammasome, and are therefore not able to assemble inflammasomes.

### Amplification of inflammasome activity is mediated by a soluble and ubiquitously produced platelet and megakaryocyte-derived factor

Phagocytosis of activated platelets has been shown to regulate platelet and neutrophil function, survival and differentiation (Badrnya et al., 2014; Chatterjee et al., 2015; Lang et al., 2002; Senzel and Chang, 2013). To determine whether phagocytosis is responsible for the effect of platelets on inflammasome activation, we performed confocal imaging of hMDMs (**Figure 4D**) and purified blood neutrophils (**Figure S2D**) incubated with freshly isolated platelets. Co-cultures were stimulated with LPS followed by inflammasome activation with nigericin. Both human macrophages (**Figure 4D**) and neutrophils (**Figure S2D**) phagocytosed platelets; however, pretreatment with the phagocytosis inhibitor Cytochalasin D did not prevent the platelet-mediated boost of IL-1β production by hMDMs (**Figure 4E**). Furthermore, platelets were able to boost the IL-1β responses of hMDMs in both direct and trans-well co-cultures (**Figure 4F**), and cell-free supernatant from resting, LPS-stimulated, or Thrombin-activated platelets also enhanced IL-1β production from hMDMs (**Figure 4G**). To exclude a direct effect of thrombin on macrophages, we stimulated BMDMs or hMDMs with thrombin in the absence of platelets (**Figure S2E**) and observed no change in IL-1β production.

Thus, the platelet influence on IL-1β production by inflammasome-activated macrophages is independent of cell contact. In contrast, the transfer of platelet supernatants to cultures of human neutrophils or monocytes did not influence the production of IL-1β (**Figure S2f**, and data not shown), suggesting that cell contact may be required for the platelet boosting effects in some cell types.

We observed that supernatants from quiescent and activated platelets had similar effects on hMDM inflammasome activation (**Figure 4G**). To determine whether this capacity of quiescent platelets was due to undetectable activation during isolation, we transferred cell-free supernatants from unstimulated human megakaryocytes to cultures of hMDMs. Similar to resting platelets, supernatants from unstimulated megakaryocytes boosted the inflammasome activity in hMDMs (**Figure 4H**). This finding points towards the presence of a soluble and ubiquitously expressed factor secreted by both platelets and megakaryocytes, which modulates the inflammasome activity of macrophages.

### Platelets license inflammasome activation on human macrophages through transcriptional regulation of NLRP3 and pro-IL-1**β**

In most immune cells, the activation of NLRP3 requires a “priming” stimulus, such as the engagement of pattern recognition receptors (PRRs) which initiate the transcription of *NLRP3*. The amount of NLRP3 available in the cytosol is a key limiting factor regulating its activation. We therefore investigated if the platelet effect on macrophages occurs at the transcriptional level during priming, or afterward during activation of the NLRP3 inflammasome. To this end, we co-cultured NLRP3-overexpressing immortalized mouse macrophages (NLRP3FiMøs) with platelets and directly stimulated these cells with Nigericin. As the LPS-priming signal is also required for the upregulation of pro-IL-1β, we assessed caspase-1 activity using a specific caspase-1 fluorescent substrate as a readout. Addition of platelets to NLRP3FiMøs did not enhance their caspase-1 activity (**Figure 5A**). These data indicate that platelets might influence the inflammasome activation of human macrophages by enhancing the transcription of NLRP3 and licensing its activation. To determine whether platelets modulate the expression of other inflammasome molecules we evaluated human macrophages cultivated alone or in the presence of platelet supernatants, which were used to minimize the detection of transcripts arising from platelets. The addition of supernatants from unstimulated platelets to macrophages boosted their expression of IL-1β (**Figure 5B**). Surprisingly, platelet supernatant was as effective as LPS at inducing the expression of NLRP3 and pro-IL-1β protein (**Figure S2B**, compare lanes 2 and 4). However, platelet supernatant did not alter the expression of pro-caspase-1 or ASC (**Figure 5B**). Importantly, exposure to platelets resulted in a 50% increase in formation of ASC specks hMDMs (**Figure 5C**), while hMDMs stained with isotype-matched IgG controls as well as platelets cultured alone did not show any ASC specks (**Figure 5C** and **S6A**). Co-culture with platelets or exposure to platelet supernatants resulted in increased cleaved caspase-1 and maturation of IL-1β (**Figure 5D-F**) in cell-free supernatants of activated hMDMs (**Figure 5D-F**), proteins that were not present in the isolated platelets (**Figure 5D-E**, PTLs alone). We also detected pro-IL-1β in cell-free supernatants of LPS-primed hMDMs, likely due to cell death caused by the high LPS dose (200ng/mL) used for priming. Furthermore, despite lacking pro-IL-1β transcripts (**Figure S4D**) and protein (**Figure 5D**, lane 11 and **Figure S5G**), platelets and their supernatants increased the concentrations of pro-IL-1β even on unstimulated hMDMs (**Figure 5D**, lanes 3-4). Altogether, these findings indicate that platelets boost inflammasome activation by enhancing the transcription of pro-IL-1β and NLRP3 in human macrophages.

**Figure 5.**
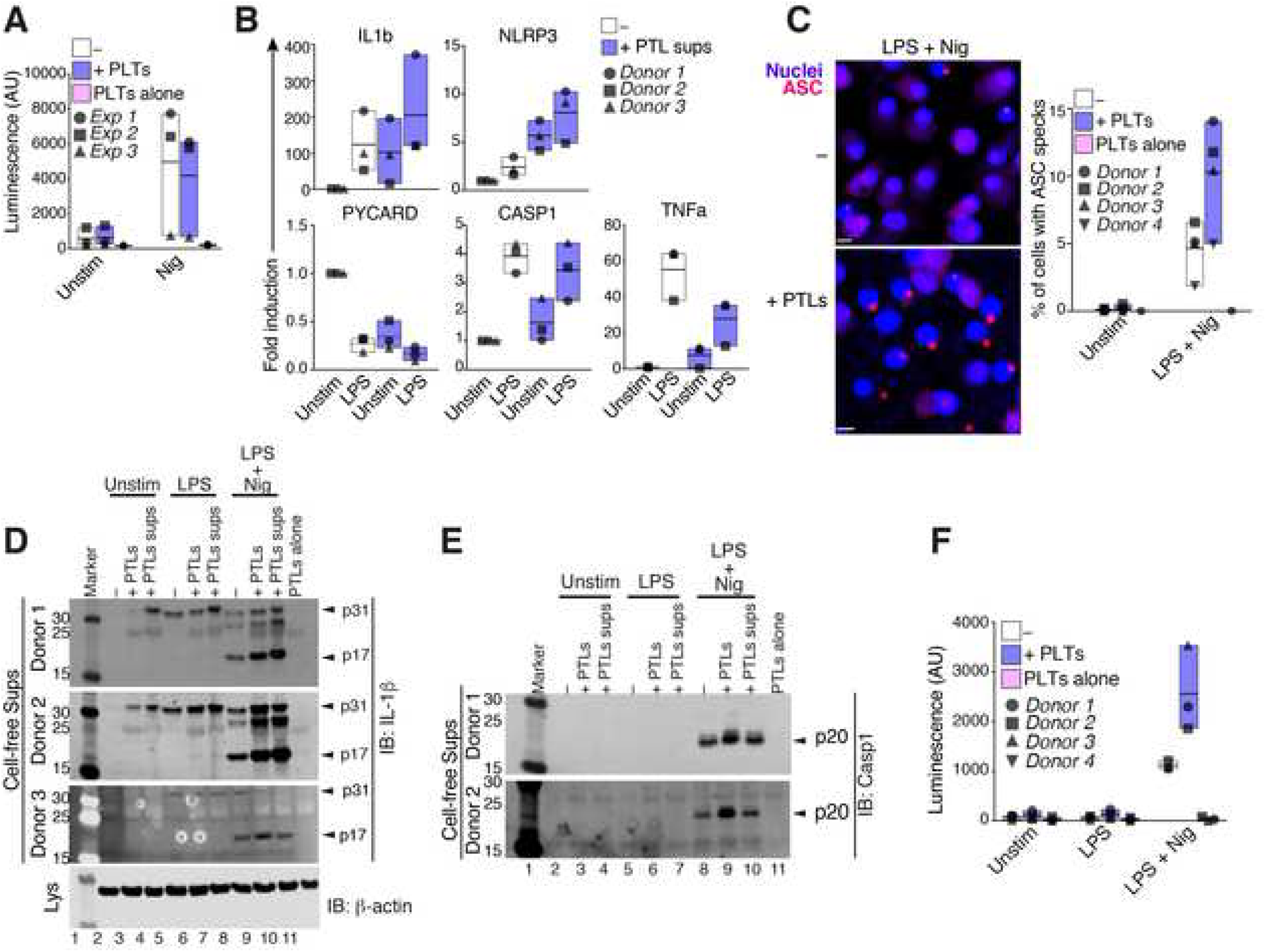
Platelets enhance NLRP3 and pro-IL-1 transcription in human macrophages. (a) Caspase-1 reporter luciferase activity of the luminogenic caspase-1 specific substrate, Z-WEHD-amino luciferin in cell-free supernatants of NLRP3-overexpressing immortalized mouse macrophages. Cells were left unstimulated (Unstim) or activated with nigericin (10µM, 90 min) without LPS priming. Cells were cultivated alone (−) or in the presence of platelets (50:1 ratio). (**B**) Real time PCR analysis of the expression of the indicated genes in human macrophages that were cultured alone, or in the presence of platelet supernatants (50:1 ratio). Cells were left untreated or stimulated with LPS. Floating bars (with mean and minimum to maximum values). Each symbol represents the average of technical triplicates from different donors (n= 3). (**C**) Representative confocal microscopy images and image-based quantification of ASC specks in LPS-primed and Nigericin-activated hMDMs that were either cultured alone (—) or in the presence of platelets (50:1 ratio). (**D**) Caspase-1 reporter luciferase activity of hMDMs treated as in c. (**E**) Immunoblotting for IL-1β, Caspase-1 in cell-free supernatants, as well as β-actin in whole cell lysates of unstimulated (Unstim), or LPS-primed hMDMs cultured alone, or in the presence of platelets, or conditioned medium from unstimulated platelets. Data from - 3 different donors are shown. (**F**) Caspase-1 reporter luciferase activity in cell-free supernatants of hMDMs treated as in c. Data in **B**, **C** and **E** is represented as floating bars (with mean and minimum to maximum values) from pooled data from 3 - 4 independent experiments. Each symbol represents the mean of technical triplicates from different donors.

### The platelet-mediated amplification of inflammasome activity requires calcium and can be prevented by heat-inactivation

We next sought to identify the platelet-derived factor that promotes macrophage inflammasome activity. Platelets are rich sources of inflammatory lipid mediators (Hinz et al., 2016). Among them, prostaglandins—COX1/2 derived mediators—have been shown to regulate LPS-induced pro-IL-1β transcription (Zasłona et al., 2017), and to inhibit TNFα in macrophages (Chandra et al., 1995; Xiang et al., 2013; Zasłona et al., 2017). We thus tested the requirement of COX1/2-derived lipid mediators for the platelet effect by inhibiting COX1/2 with Aspirin (Xiang et al., 2013). Aspirin pre-treatment of platelets did not influence on their effects on macrophage IL-1β production (**Figure 6A**). Similarly, pre-treatment of platelets with zileuton (Zt), an inhibitor of the 5-lipoxygenase enzyme involved in leukotriene synthesis, did not prevent the platelet effect. Furthermore, neither Aspirin nor Zileuton affected macrophage IL-1β production hMDMs when added directly in the absence of platelets (**Figure 6A**). These experiments indicate that the effect of platelets on the macrophage inflammasomes is independent of COX1/2-derived lipid mediators and 5-LOX-induced leukotrienes.

**Figure 6.**
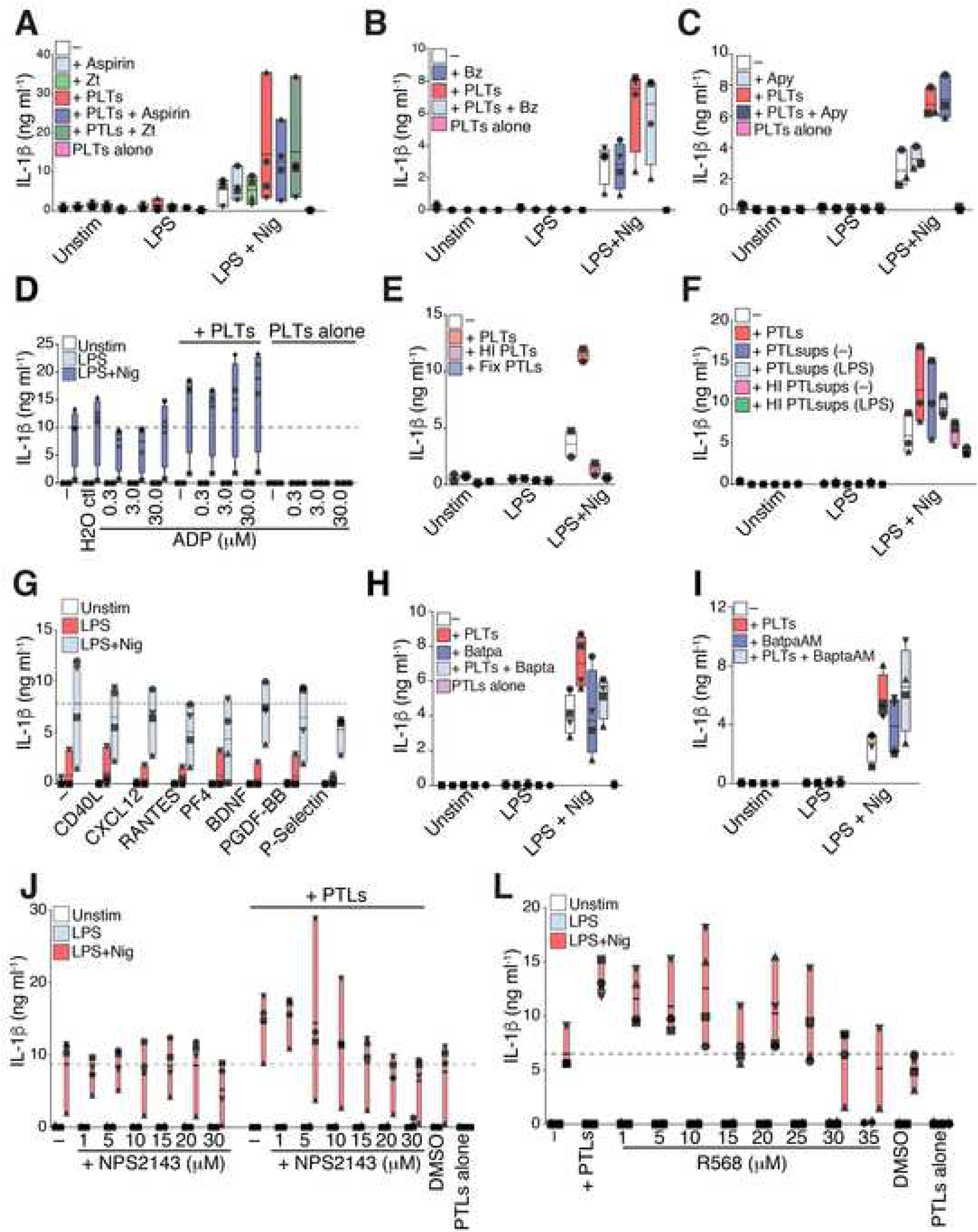
A platelet-derived calcium-dependent protein boost the inflammasome activity of human macrophages. (a) IL-1β levels in cell-free supernatants of unstimulated (Unstim), or LPS-primed (200 ng/mL, 3 hours), and nigericin-activated (10 µM, 90 min) human monocyte-derived macrophages (hMDMs). Cells were cultivated alone (−), or in the presence of platelets (+PTLs, 50:1 ratio) that were pre-treated with Aspirin (100 µM), or Zileuton (Zt, 100 µM) for 1 hour before being added to macrophages (n. = 4). (**B**) IL-1β levels measured in cell-free supernatants of unstimulated, or LPS-primed hMDMs that were added with platelets. Co-cultures were incubated with BenzonaseR (**B**) or Apyrase 0.5 U/ml (**C**) before LPS priming and inflammasome activation with nigericin (n. = 4). (**D**) IL-1β levels in cell-free supernatants of co-cultures of platelets and hMDMs added with the indicated concentrations of ADP or equivalent dilutions in water (n. = 4). (**E-F**) IL-1 β levels in cell-free supernatants of hMDMs stimulated as in **A**, and cultured alone or in the presence of heat-inactivated (HI, 80°C for 40 min), of cross-linked (4% paraformaldehyde) platelets (n = 2), or (**D**) platelet supernatants (n = 3). (**G**) IL-1β levels in cell-free supernatants of hMDMs activated as in a, and added with the indicated recombinant human proteins (n. = 3-4). (**H**) IL-1β levels in cell-free supernatants of co-cultures of platelets and hMDMs added with BAPTA (0.5 mM) in calcium-free medium or (**I**) BAPTA AM (5 μM) in normal RPMI medium before nigericin stimulation (n. = 4). (**J**) IL-1β levels in cell-free supernatants of inflammasome-activated hMDMs incubated with the indicated concentrations of calcium chloride (CaCl_2_) in Ca^2+-^free medium (n. = 3). Data is represented as floating bars (with mean and minimum to maximum values) from pooled data from 2 - 4 independent experiments. Each symbol represents the mean of technical triplicates from different donors. (See also **Figure S6**).

Activated platelets can also release nucleic acids and nucleosides which display inflammatory properties (Qin et al., 2016). However, degradation of free nucleic acids by addition of BenzonazeR did not prevent the effect of platelet co-culture on IL-1β production, indicating the effect is not mediated by platelet-derived nucleic acids (**Figure 6B**). As platelets are rich stores of other purines including ADP and ATP, a *bona fide* activator of NRLP3, we speculated that platelet-released ATP could boost macrophage inflammasome activity. To address this, we added apyrase, which hydrolyzes both ATP and ADP, to the co-culture of hMDMs with platelets. Apyrase addition did not alter the effect of platelets on hMDM inflammasome activity (**Figure 6C**). Furthermore, direct addition of extracellular ADP to inflammasome-activated hMDMs cultivated alone or together with platelets did not affect IL-1β production (**Figure 6D**), indicating that platelet-derived ATP and ADP do not play a role in this context. This is in line with our findings that platelets boost macrophage IL-1β production through transcriptional regulation during the priming phase of inflammasome activation (**Figure 5A-B**).

Therefore, we speculated that the platelet-derived factor may be a protein. In line with this hypothesis, both heat inactivation and crosslinking of platelets (**Figure 6E**) and platelet supernatants (**Figure 6F**) extinguished the effect on inflammasome activation in human macrophages. Multiple cytokines were detectable in cell-free supernatants of resting and activated platelets in our Luminex assays (**Figure S1A**). Those included: CCL5 (RANTES), CXCL12 (SDF1α), IL-18, and PDGF-BB. Additional literature reported that platelets are cellular sources of CD40L, PF4, BDNF, P-selectin, and CXCL7 (Kral et al., 2016; Semple et al., 2011). To evaluate the potential roles of these factors in boosting macrophage inflammasome activity, we employed several experimental approaches. Those included: i) direct addition of human recombinant proteins to hMDMs (**Figure 6G**, **Figure S6B-C**); ii) addition of inhibitors or blocking antibodies against human platelet-derived cytokines to co-cultures of hMDMs and platelets (**Figure S6D-E**); and iii) experimental co-cultures involving cells from animals deficient for chemokine receptors (**Figure S6F**). Altogether, these experiments excluded a role for CXCL1, CCL5, CXCL12, CXCL7, PDGF-BB, EGF, VEGFA, CD40L, PF4, BNDF and P-selectin as contributors to the platelet effect.

The failure of the above mentioned α-granule-derived proteins to boost the IL-1β production of human macrophages suggested that molecules contained in the dense granules of platelets could play a role in the platelet-mediated regulation of IL-1β in macrophages. Platelet dense granules are rich stores of ADP, ATP, serotonin (5-HT), and Ca^2+^. We therefore tested whether platelet-derived 5-HT could influence IL-1β production in macrophages. We found that neither the addition of recombinant 5-HT (**Figure S6G**) nor the blockade of its uptake by macrophages with the selective 5-HT reuptake inhibitor fluoxetine (Du et al., 2016; Marcinkiewcz et al., 2016) (**Figure S6H**) altered IL-1β production by inflammasome-activated hMDMs.

Next, we examined the effect of chelating extracellular and intracellular calcium with BAPTA and BAPTA-AM, respectively. Notably, treatment of co-cultures with BAPTA, but not with the membrane permeable BAPTA-AM, prevented the platelet effect without interfering with the standard response of hMDMs to LPS + Nigericin (**Figure 6H-I**). However, when added directly to hMDMs cultured without platelets, calcium chloride (CaCl_2_) did not evoke additional IL-1β secretion from inflammasome-activated hMDMs (**Figure S6**), suggesting that Ca^2+^ is required but not sufficient for the platelet effect.

Cells sense extracellular Ca^2+^ through two G-protein-coupled receptors (GPCRs), CaSR and GPRC6A. The requirement of Calcium for the platelet effect on human macrophages suggests that a calcium sensing receptor (CaSR) or G-coupled receptors might be involved. To test this hypothesis, we pre-treated hMDMs with two selective inhibitors of CaSR, NPS2143 and Calhex231. We found that inhibition of CaSR with NPS2143 (**Figure 6J**) but not with Calhex231 (data not shown), blocked the platelet-mediated effect on hMDMs. Of note, similar to platelets, allosteric activation of CaSR with R-568 boosted the IL-1β response of inflammasome-acrtivated human macrophages (**Figure 6L**). These findings indicate that extracellular Ca2+ is involved on the platelet-mediated boosting of IL-1β of inflammasome-activated human macrophages. Extracellular Ca2+ has been previously proposed to activate the NLRP3 inflammasome (Rossol et al., 1AD). Taken together, these findings indicate that a heat-sensitive protein that induces calcium signaling is involved in the regulation of the inflammasome activity of human macrophages.

### A combination of platelet-derived proteins might boost inflammasome activation of innate immune cells

Next, we employed a proteomics approach to assess the secretome of platelets and megakaryocytes to identify additional proteins similarly secreted by these cells that could mediate the IL-1 boosting effect (**Figure 7A-B**, and **Figure S7A-B**). For this, cell-free supernatants from quiescent, or LPS-treated human platelets (**Figure. 7A**, n = 4) as well as from the human megakaryocytic cell line MEG-01 (**Figure 7B**, n = 3) were assessed by Mass Spec proteomics, using the label-free quantification (LFQ) method. As our findings on **Figure 4G-H** indicated the presence of a factor ubiquitously present in the supernatants of resting and activated platelets, we directed further analysis to proteins that remained unchanged by LPS stimulation in both cell types (fold change >-1.5 and < 1.5). Among the proteins similarly secreted by platelet and megakaryocytes, we identified several members of the TGF-β family of proteins, S100 proteins and thrombospondin-1 (THBS1), which Figure d as the highest abundant protein in platelet supernatants (**Figure 7A-B**). These findings were in line with the platelet-induced transcriptional changes on hMDMs. Notably, TGF-β, S100 and THSB1 require calcium for their activity (Bertheloot and Latz, 2017a; Cailotto et al., 2011; Misenheimer and Mosher, 1995). Additionally, THBS1 was also shown to activate TLR4 (Li et al., 2013) and regulate IL-1β of macrophages (Stein et al., 2016), and to be necessary for the full activity of TGF-β in vivo (Crawford et al., 1998).

**Figure 7.**
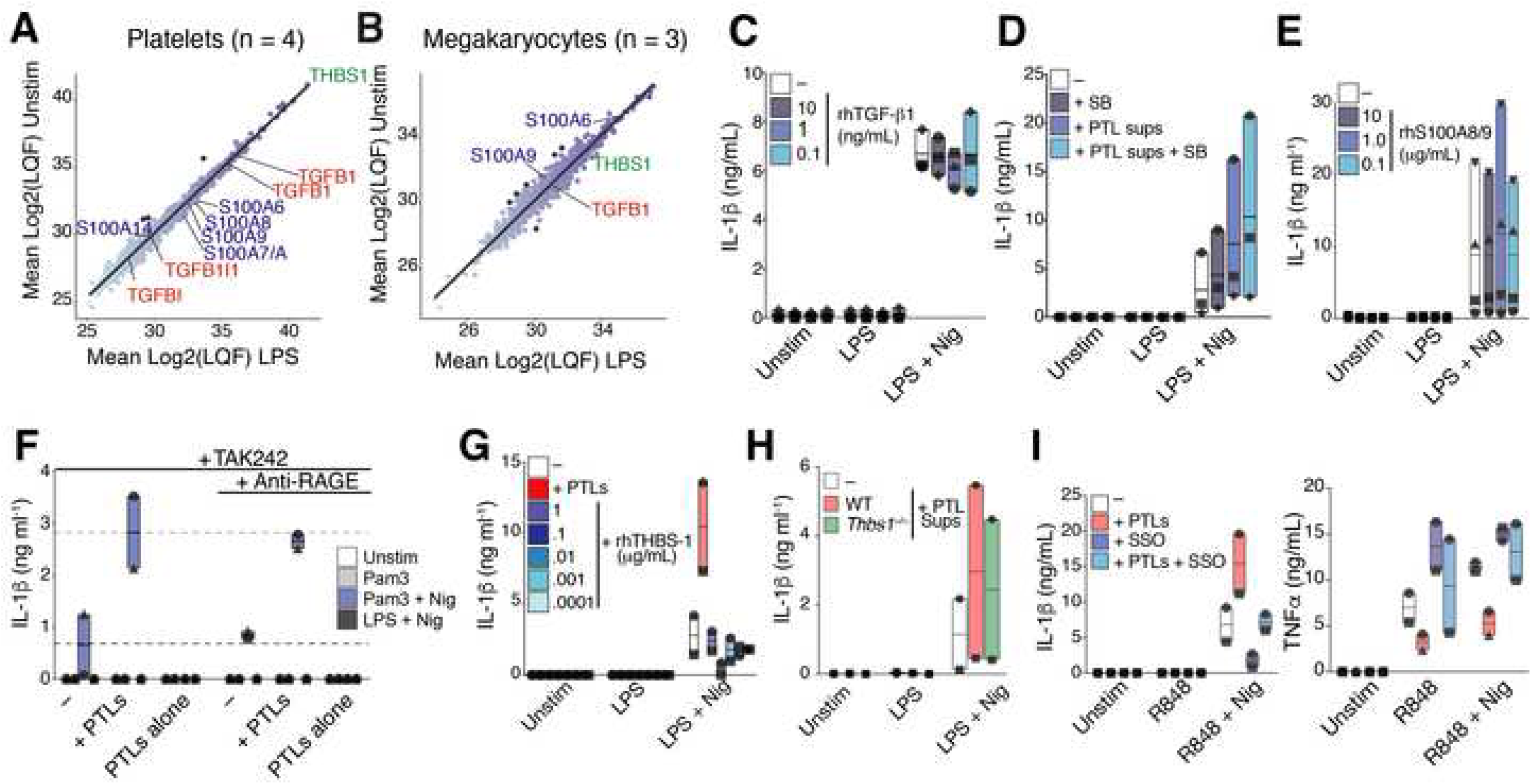
Platelets have broad effects on macrophages that may contribute to boosting of inflammasome activation. (**A - B**) Label-free quantification (LFQ) proteomic assessment of the proteins in cell-free supernatants from unstimulated (Unstim) or LPS-treated (**A**) human platelets (n = 4), or (**B**) the human megakaryocytic cell line MEG-01 (n = 3). Dark-blue points show proteins with Log2 fold higher or lower than 1.5 between LPS and unstimulated conditions. (**C - E**) IL-1β levels in cell-free supernatants of unstimulated or inflammasome-activated hMDMs that were (**C**) added with recombinant human TGFb1 in the indicated concentrations (n = 3), or (**D**) pre-treated with SB-431542 (10 µM, for 1 hours) (n = 3). (**E - F**) IL-1β levels in cell-free supernatants of unstimulated or inflammasome-activated hMDMs that were (**E**) added with recombinant human S100A8/9 (n = 4), or (**F**) pre-treated with TAK242 (0.5 µg ml^-1^), and an Anti-RAGE mAb (10 µg ml^-1^), n = 2. (**G**) IL-1β levels in cell-free supernatants of unstimulated or inflammasome-activated hMDMs that were added with the indicated concentrations of recombinant human Thrombospondin-1 (n = 2). (**H**) Levels of IL-1β in cell-free supernatants inflammasome-activated mouse BMDMs cultured alone or in the presence of supernatants isolated from wild-type or *Thbs1*^−/−^ platelets. (**I**) IL-1β and TNFα levels in cell-free supernatants of unstimulated or R848-primed and Nigericin-activated hMDMs cultured alone or in the presence of platelets (50:1). hMDMs were pre-treated with the CD36 inhibitor Sulfosuccinimidyl oleate (SSO, n = 2). **C** –**I** Floating bars (with mean and minimum to maximum values) from pooled data from 2-4 independent experiments. Each symbol represents different donors, or mice. (See also **Figure S7**).

We therefore tested the effects of these proteins on the platelet-mediated boosting of inflammasomes. To evaluate the role of TGF-β, we tested the addition of recombinant human TGF-b1 to inflammasome-activated hMDMs. Consistent with previous reports of an antagonistic effect of TGF-β on TNFα signaling (Vaday et al., 2001; Verrecchia and Mauviel, 2004; Yamane et al., 2003), addition of recombinant human TGF-β1 (rhTGF-β) to hMDMs inhibited their production of TNFα triggered by LPS (**Figure S7C**). However, rhTGF-β did not influence the IL-1β response (**Figure 7C**). Likewise, selective inhibition of the TGF-β receptor by pre-treating hMDMs with the small molecule SB431542 (Matsuyama et al., 2003) did not prevent the platelet effect (**Figure 7D**), despite preventing down-regulation of TNFα (**Figure S7C**). These data indicate that, TGF-β is likely not the platelet-derived factor enhancing the inflammasome activity of human macrophages, but might underlie the regulation of TNFα.

S100 proteins are important regulators of calcium homeostasis, and platelet-derived S100A8/9 has been shown to induce inflammation in several inflammatory diseases (Lood et al., 2016; Wang et al., 2014). S100A8/9 is enriched in platelets and binds to various membrane receptors such as TLR4, RAGE, and CD36 (Bertheloot and Latz, 2017b). TLR4 and CD36 are also known receptors for THBS1 (Li et al., 2013; Stein et al., 2016). However, we found that neither addition of recombinant human rhTHBS1 (**Figure 7G**) nor rhS100A8/9 (**Figure 7E**) to human macrophages influenced their basal response to LPS + Nigericin. Similarly, cell-free supernatants from platelets from *Thbs1*^−/−^ mice were as efficient as WT platelets in boosting IL-1β release, caspase-1 activity from inflammasome-activated BMDMs (**Figure 7H**). Of note rhS100A8/9 induced TNFα secretion from hMDMs (**Figure S7D**). Similarly, inhibition of RAGE using a specific anti-RAGE antibody revealed no effect on the platelet boosting of IL-1β release (**Figure 7F** and **S7G**). These findings led us to conclude that platelet-derived THBS1, though abundantly secreted by platelets and megakaryocytes, is not responsible for the platelet effect on macrophages.

As THBS1 and S100 proteins bind to CD36, we tested the effect of CD36 inhibition using Sulfosuccinimidyl oleate (SSO) (Kuda et al., 2013). SSO interfered with IL-1β and TNFa release (**Figure 7I**) and Caspase-1 activity (**Figure S7F**) both in the presence and absence of platelets. Nevertheless, even in the presence of SSO, platelets increased both the release of IL-1β and the activity of Caspase-1 from macrophages compared to SSO pre-treated macrophages in monoculture.

As single inhibition of TLR4, RAGE, and CD36 did not completely extinguish the platelet effect, we speculated that a combined activity of platelet-derived S100 and THBS1 could mediate the boosting of IL-1β on hMDMs. An experimental strategy to block TLR4/RAGE and CD36 resulted in complete inhibition of IL-1β, TNFα and Caspase-1 activity of human macrophages (**Figure S7G**) thus precluding us to conclude whether these proteins are synergistically involved in the platelet-mediated enhancement of IL-1β response of human macrophages. Hence, additional work will be required to precisely delineate mechanisms by which platelets affect the inflammasome activation of human macrophages.

## DISCUSSION

In summary, we report that interaction with platelets licenses NLRP3 and potentiates inflammasome activation and production of IL-1 cytokines by innate immune cells. We show that the platelet effect is mediated by a soluble protein factor that engages CaSR on macrophages. Our findings reveal a novel role for platelets in governing the production of IL-1 cytokines and emphasize that the regulation of the inflammasome *in vivo* is too complex to be modeled by *in vitro* monocultures.

Our work highlights the previously underappreciated function of platelets in modulating immunity, adding to the growing evidence that platelet parameters, such as counts and activation, are causes of variation in the concentrations of circulating cytokines (Hu et al., 2018; Schirmer et al., 2018; Tunjungputri et al., 2018). Although best known for their role in thrombosis, platelets secrete a host of proteins (Coppinger, 2004; Maynard et al., 2007), and can modulate the function of leukocytes by forming heterotypic aggregates (Kral et al., 2016). The interaction of platelets with macrophages has been less explored, in part due to the assumption that platelets have limited access to tissues, where most macrophages reside. We show here that the effects of platelets on macrophages, but not on neutrophils, is independent of cell contacts, which could overcome the tissue barriers separating these cells. However, recent discoveries have shown that platelets can reach deeply into areas of tissue that are not readily accessible by other immune cells, such as the core of a tumor (Best et al., 2015, 2017). Platelets were found in the inflammatory tissue site, in where their exert critical functions in bacterial clearance, hours before the arrival of the first neutrophils (Wong et al., 2013). Moreover, platelets guide neutrophils and promote their extravasation from blood into tissue (Alard et al., 2015; Asaduzzaman et al., 2009; Sreeramkumar et al., 2014; 2017; Zuchtriegel et al., 2016).

Our work also challenges the role of inflammasomes on platelets. Although a few reports describe the assembly of inflammasomes on platelets from patients with Dengue fever (Hottz et al., 2013), sickle cell disease (Vogel et al., 2018), and sepsis (Cornelius et al., 2019), we used multiple complementary techniques to demonstrate that platelets lack expression of NLRP3, ASC, and Caspase-1, and are therefore incapable of assembling inflammasomes. This conclusion was further supported by a meta-analysis of publicly available human platelet transcriptome data from 5 independent studies. The discrepancy with previous results may be because other studies relied on antibody-based fluorescence assessment of inflammasome components on platelets, without quantitative measurement of the relevant proteins in platelet lysates. Furthermore, most of the previous studies employed antibodies with weak specificity, which have been shown to react with ASC-deficient cells (Beilharz et al., 2016).

Although the expression of IL-1 cytokines in platelets has also been reported, the results are conflicting, with some studies describing expression of IL-1 in activated platelets (Denis et al., 2005; Lindemann et al., 2001; Thornton et al., 2010b), and others finding IL-1 activity in bioassays (Hawrylowicz et al., 1989; Kaplanski et al., 1993), without evidence of the cytokine itself. Platelets have been reported to amplify IL-1β-mediated inflammation in a rheumatoid arthritis (RA) model (Boilard et al., 2010), however, it was not possible in that study to distinguish whether the enhanced IL-1β response was caused by platelet-derived IL-1β or their influence on other immune cells as we demonstrated here. Importantly, the expression of IL-1β on platelets has been closely correlated to the presence of contaminating leukocytes in platelet preparations (Pillitteri et al., 2007). We observed that platelets are poor producers of IL-1β in all the experimental conditions tested (priming with TLR2, TLR4, TLR7/8 ligands followed by inflammasome activation with Nigericin, ATP, or PrgI). Notably, IL-1β expression in platelets has been demonstrated after 18 hours of thrombin stimulation *in vitro*, (in the range of 150 pg. ml^-1^) (Brown and McIntyre, 2011), however, we detected robust IL-1β production in macrophage-platelet co-culture experiments at values greater than >6000 pg. ml-1 in 4.5 hours, indicating that the possible contribution of platelet-derived IL-1β was minimal. Finally, our experiments in cells from mice deficient in IL-1β, IL-1R, or inflammasome components exclude a role for platelet-derived inflammasomes or IL-1β in the platelet effect.

Our data suggest a role for platelets in regulating the production of inflammasome-derived IL-1 cytokines that may be relevant to human physiology and disease. In mice, we found that platelet depletion decreased serum levels of IL-1β, though not significantly. The regulation of IL-1β *in vivo* is complex; although it is induced within 15 minutes following LPS stimulation, it rapidly declines due to a short mRNA half-life and other mechanisms (Dinarello, 2009), which could underlie our difficulty in capturing the effect. However, multiple studies support a role for platelets in modulating human inflammatory diseases. High blood platelet counts are found in patients (Ciccarelli et al., 2014) and mouse models of IL-1-driven auto-inflammatory disorders (Bonar et al., 2012). In Kawasaki Disease (KD), an acute systemic vasculitis in children for which thrombocytosis is a common feature, recent human and experimental mouse models have determined a critical role of IL-1β in the cardiovascular pathogenesis (Burns et al., 2017; Lee et al., 2012b). Indeed, the degrees of thrombocytosis and platelet activation are predictive of the development of coronary aneurysm in KD.

As both IL-1α and IL-1β induce thrombocytosis in mice (Kimura et al., 1990; Nishimura et al., 2015; Trinh et al., 2015), our findings raise the possibility that platelets could help feed an inflammatory loop by potentiating and sustaining IL-1 signaling, which in turn stimulates platelet biogenesis. However, whether thrombocytosis is a marker of disease or a contributor to pathogenesis, remains to be investigated.

Platelet counts have also been correlated with the concentrations of other cytokines that affect platelet production. For example, low platelet counts are associated with lower levels of CD40L, CXCL5, CCL5, and EGF in the plasma of patients with aplastic anemia and immune thrombocytopenic purpura (Feng et al., 2012), two diseases characterized by lower blood platelet counts. Another study reported higher levels of VEGF, GM-CSF, IFNγ, MCP-1, IL-8, and PDGF-BB strongly associated with higher platelet counts in patients with essential thrombocythemia, compared to polycythemia vera (Pourcelot et al., 2014). Thus, the contribution of platelets to cytokine production in human biology warrants further investigation.

Notably, our results also highlight a role for platelets in modulating TNFα production from macrophages. In agreement with this finding, a recent study reported a protective role of platelets in sepsis through the regulation of TNFα and IL-6, via the secretion of a COX1/PGE2/EP4 dependent lipid mediator (Xiang et al., 2013). We observed that TNFα is regulated via a platelet-derived COX1-dependent lipid mediator, and related to TGFβ signaling.

Despite our efforts, we were not able to identify a unique platelet-derived factor that drives the effect of platelets on innate immune cell inflammasomes. However, we show that the regulation of IL-1β production is likely mediated through a constitutively expressed protein factor that is sensitive to calcium. We also observed that platelets trigger calcium sensing receptors (CaSR) on macrophages, consistent previous report of a role for CaSR on inflammasome activation(Lee et al., 2012a). However, these findings will need further validation in genetic deficiency of these receptors. Because a combination of factors acting in synergy and affecting different pathways may underlie the effects we observed, refined experimentation will be necessary to elucidate and validate the molecular mechanisms.

Our findings also add to recent new discoveries demonstrating direct links between the mammalian immune and coagulation system (Burzynski et al., 2019; Wu et al., 2019; Zhang et al., 2019). For instance, it was found that pro-IL-1α is cleaved and processed by thrombin at a perfectly conserved site across disparate species (Burzynski et al., 2019) and that coagulation mediates both host defense(Zhang et al., 2019) and lethality caused by overactivation of inflammasomes (Wu et al., 2019). Our study strengthens this link by showing a direct participation of platelets in the inflammasome potential of innate immune cells.

## Acknowledgments

We thank Feng Shao (National Institute of Biological Sciences, Beijing 102206, China), for the plasmid encoding the PrgI-protective antigen conjugate protein, as well as Matthias Geyer and David Fußhöller for the purified PrgI protein used here to activate the NLRC4 inflammasome. We are grateful to Damien Bertheloot for critically reading our manuscript, and to Matthew S. Mangan, as well as members of the Institute of Innate Immunity for helpful discussions. This study was supported by a grant from the European Research Council (PLAT-IL-1). B.S. Franklin is supported by grants from the German Research Foundation (DFG, SFBTRR57). B.S. Franklin and E. Latz are members of the ImmunoSensation cluster of Excellence in Bonn. E. Latz is supported by grants from the DFG (SFB645,704, 670, 1123, TRR57, 83) and the European Research Council (InflammAct). V. Rolfes is supported by a fellowship from Bayer.

## Author contributions

Conceptualization, B.S.F.; Investigation, B.S.F., V.R., I.H., L.S.R., L.B., N.R., S.M., M.L.S.S., M.R., & H.J.S.; Data Analysis, B.S.F., & S.V.S.; Resources, B.S.F., M.A., C.J.F. and E.L.; Software, B.S.F. and S.V.S.; Writing – Original Draft, B.S.F.; Revisions, M.A., V.R. and L.S.R.; Visualization, A.M, V.R., L.S.R., and I.H.; Supervision, B.S.F., L.H.C.; Project Administration, B.S.F..; Funding Acquisition, B.S.F.

## Declaration of Interests

“The authors declare no competing interests.”

## METHODS

### Patients and human volunteers

Malaria patients naturally infected with *Plasmodium vivax* in the Amazon area of Cuiaba (Mato Grosso, Brazil) were invited to participate in the study. Seventy-eight individuals (aged18 - 78 years old) who sought care at the Julio Muller Hospital, whose thick blood smear was positive for *P. vivax* were included. Another 9 healthy volunteers from the same endemic area who tested negative for Plasmodium infection were recruited and served as healthy donor controls. Exclusion criteria included: (i) refuse or inability to sign the informed consent; (ii) age <18 years; (ii) pregnant women; (ii) mixed infection with *P. falciparum* or *P. malaria*, tested by both microscopic examination and a nested-PCR; (iv) any other co-morbidity that could be traced. Clinical and demographical data were acquired through a standardized questionnaire, and the hematological profiles were assessed by automated complete blood count carried out at the site hematology facility. Plasma samples were isolated immediately after blood sampling and stored at −80°C until use. The study was approved by the Ethical Review Board of the René Rachou Research Center, FIOCRUZ, Brazilian Ministry of Health (Reporter CEPSH/CPqRR N. 05/2008 and N. 01/2018).

All participants were instructed about the objectives of the study and signed an informed consent in accordance with guidelines for human research, as specified by the Brazilian National Council of Health (Resolution196/96). Patients diagnosed with *P. vivax* malaria were treated according to the standard protocols recommended by the National Malaria Control Program (chloroquine + primaquine).

### Generation of human primary macrophages (hMDMs)

Buffy coats from healthy donors were obtained according to protocols accepted by the institutional review board at the University of Bonn (local ethics votes Lfd. Nr. 075/14). Primary human macrophages were obtained through differentiation of CD14+ monocytes in a medium complemented with 500 U/mL rhGM-CSF (Immunotools) for 3 days. In brief, human peripheral blood mononuclear cells (PBMCs) were obtained from buffy coats of healthy donors by density gradient centrifugation in Ficoll-Paque PLUS (Healthcare). PBMCs were incubated at 4°C with magnetic microbeads conjugated to monoclonal anti-human CD14 antibodies according to the manufacturer’s instructions (Miltenyi Biotech). CD14+ monocytes were thereby magnetically labeled and isolated using a MACS column placed in a magnetic field. CD14+monocytes were cultivated in complete medium (RPMI1640 medium with 10% FBS, 1% Penicillin-Streptomycin, 1% GlutaMAX and 1% Sodium Pyruvate) complemented with 500 U/mL rhGM-CSF at a concentration of 2×10^6^/mL in 6-well plates to generate monocyte-derived macrophages. Cells were harvested at day 3, counted using a hemocytometer and seeded at a concentration of 1×10^5^/well in complete medium complemented with 125 U/mL rhGM-CSF in 96-well flat-bottom plates and incubated overnight for experiments on the next day.

### Generation of murine bone marrow-derived macrophages (BMDMs)

Mice were anaesthetized with isoflurane and sacrificed by cervical dislocation. Femur and tibia from hind limbs were removed and the bones were briefly disinfected with 70% ethanol. The bone marrow cavity was flushed with PBS and the cell suspension was filtered through a 70 µm cell strainer before centrifugation at 400 × g for 5 minutes. Cells were resuspended in DMEM supplemented with 20% L929 supernatant and cultured for 6 days to differentiate into macrophages (BMDMs). On day six, cells were harvested using cold PBS containing 5 mM EDTA and 2% FBS and scraping. After centrifugation at 350 × g for 5 minutes, the BMDM were seeded at 1×10^5^/well in DMEM with 20% L929 supernatants in flat-bottom 96-well plates and incubated overnight for experiments on the next day.

### Human and mouse cell isolations

Peripheral blood was obtained by venipuncture of healthy volunteers after signature of informed consent, and approval of the study by the Ethics Committee of the University of Bonn (Protocol#282/17), and in accordance with the Declaration of Helsinki.

### Neutrophil isolation from human blood

Venous blood was collected in S-Monovette^®^ K3EDTA tubes and neutrophils were isolated using the EasySep^TM^ Direct Human Neutrophil Isolation Kit according to the manufacturer instructions (STEMCELL Technologies^TM^). Whole blood was incubated with the Neutrophil Isolation Cocktail and RapidSpheres^TM^ for 5 minutes and diluted with neutrophil isolation buffer (1mMEDTA in PBS). After 5 minutes of incubation in the EasySep^TM^ Magnet, the enriched cell suspension was poured into a new tube and incubated again with RapidSpheresTM for another 5 minutes, followed by a second, and third round of magnetic separation. The obtained neutrophils were counted using a hemocytometer and pelleted by centrifugation at 350 × g for 5 minutes. Cells were resuspended in RPMI-1640 medium supplemented with 10% FBS, 1% GlutaMAX and 1% Penicillin-Streptomycin. Neutrophil suspension was adjusted to 1×10^6^/mL and 100 µL (1×10^5^cells/well) were seeded in a 96-well round-bottom plate. The purity of the purified neutrophils was assessed by flow cytometry using CD66b (neutrophil marker) and CD41 (platelet marker).

### Platelet isolation from human blood

Human platelets were isolated as previously described (Alard et al., 2015) with slight modifications. In brief, venous blood was drawn into S-Monovette^®^ 9NC collection tubes. The blood was centrifuged for 5 minutes at 330 × g without brake to obtain platelet-rich plasma (PRP). All following centrifugation steps were performed without brake and in the presence of 200 nM PGE1 to inhibit platelet activation. PRP was transferred to a new tube and diluted 1:1 with phosphate-buffered saline (PBS) to reduce leukocyte contamination and centrifuged for 10 minutes at 240 × g. Platelets were pelleted by centrifugation at 430 × g for 15 minutes and washed once with PBS. Total platelets were counted using a hemocytometer and resuspended in RPMI medium to a concentration of 1×10^8^/mL unless otherwise indicated. The purity of the purified platelets was assessed by flow cytometry using CD45 (leukocyte) and CD41 (platelet) markers.

### Generation of Platelet and megakariocyte supernatants

After platelet isolation, the cell suspension was adjusted to 5×10^7^ platelets ml^-1^ (for human experiments) or 5×10^6^ platelets ml^-1^ (for mouse BMDM experiments), in order to fit the macrophage:platelets proportion of 1:50 (human) or 1:5 (mouse). Next, platelets were left unstimulated (Unstim) or incubated at 37° C with LPS (200 ng ml^-1^) or thrombin (0.5 or 1.0 U ml^-1^) for 3 hours. After stimulation, the cell suspension was centrifuged for 10 minutes at 3000xg for the generation of the cell-free supernatants. The absence of cells was confirmed by microscopic visualization in a hemocytometer. The supernatants were immediately used to stimulate hMDMs in RPMI, or immediately frozen at −80° C until use.

### Monocyte isolation from human blood

Venous blood was collected in S-Monovette^®^ K3EDTA tubes and PBMCs were obtained by density gradient centrifugation in Ficoll-Paque PLUS. Monocytes were isolated from PBMCs using the EasySepTM Human Monocyte Isolation Kit according to the manufacturer instructions (STEMCELL Technologies^TM^). PBMCs were washed twice with PBS complemented with 2% FBS and 1mM EDTA before incubated with the supplied monocyte isolation cocktail and the platelet removal cocktail for 5 minutes. Magnetic beads were added to this suspension for another 5 minutes before magnetic separation in an EasySepTM Magnet. After 2.5 minutes of incubation, the enriched suspension was poured into a new tube. The isolated monocytes were counted using a hemocytometer and resuspended in RPMI medium at a concentration of 1×10^6^/mL. The purity of the purified monocytes was assessed by flow cytometry using CD14 (monocyte) and CD41 (platelet) markers.

### Animals

Mice were housed under standard conditions at 22°C and a 12 h light-dark cycle with free access to food and water. Animal care and handling was performed according to the Declaration of Helsinki and approved by the local ethical committees (LANUVNRW # 84-02.04.2016.A487).

### Platelet isolation from murine blood

Blood was drawn by puncturing the vena facialis of anaesthetized mice. Blood from mice of the same genotype were pooled in a sterile 5 mL polystyrene tube containing one-sixth blood volume of pre-warmed citrate-dextrose solution (ACD). PRP was prepared by centrifugation at 330 × g for 5 minutes without brake. All following centrifugation steps were performed without brake. PRP was transferred to a new tube and diluted in twice as much volume of PIPES/saline/glucose (PSG)buffer with the final concentration of 1.5 µM PGE1. The suspension was centrifuged at 240 × g for 10 minutes to reduce leukocyte and erythrocyte contamination. The supernatant was transferred into a tube with PGE1 in a final concentration of 0.7 μM in PSG buffer. The platelets were pelleted by centrifugation at 1000 × g for 5 min, washed with 1.5 μM PGE1 in PSG buffer. The washed platelets were resuspended in DMEM, counted in a hemocytometer, and the platelet suspension was adjusted to 5×10^6^/mL unless otherwise indicated. Purity and viability of the prepared platelets were assessed by flow cytometry.

### Purity assessment of the isolated cells by flow cytometry

Samples of isolated neutrophils and platelets were analyzed for purity, platelet pre-activation and platelet viability after each experiment. Isolated murine and human platelets were activated with 0.5 or 1 U/mL thrombin respectively for 30 min. Cells were blocked with 1:10 mouse or human Fc blocking reagent for 10 minutes at room temperature (RT). The samples were stained with fluorochrome-conjugated monoclonal anti-mouse or anti-human Ig antibodies against CD41/CD41, CD62p, CD14, CD45 or CD66b as indicated for 30 minutes in the dark. Cells were washed and resuspended in flow cytometry buffer (1% FBS in PBS) for analysis. Compensation beads (OneComp Beads) and isotype controls were prepared in the same way. Flow cytometry was performed with a Macs Quant^®^ Analyzer10 (Miltenyi Biotech) and analyzed using the Flowtop software (Tree Star). The applied gating strategy was based on doublet discrimination and isotype-matched control antibodies.

### Generation of Platelet and megakaryocyte cell-free supernatants

After platelet isolation, the cell suspension was adjusted to 5×10^7^ platelets ml^-1^ (for human experiments) or 5×10^6^ platelets ml^-1^ (for mouse BMDM experiments), in order to fit the macrophage:platelets proportion of 1:50 (human) or 1:5 (mouse). For the generation of supernatants from the megakaryocytic cell line MEG-01, 1×10^6^ cells ml^-1^ were used. Next, platelets or MEG-01 were left unstimulated (Unstim) or incubated at 37° C with LPS (200 ng ml^-1^) or thrombin (1.0 U ml^-1^) for 3 hours. After stimulation, the platelet cell suspension was centrifuged for 10 minutes at 3000 × g for the generation of the cell-free supernatants. MEG-01 cells were first centrifuged for 7 minutes at 170 × g and the obtained supernatants were centrifuged again for 10 minutes at 3000 × g. The absence of cells was confirmed by microscopic visualization in a hemocytometer. The supernatants were immediately used to stimulate hMDMs in RPMI, or immediately frozen at −80° C until further use.

### Stimulation assays

Seeded BMDMs, human macrophages or neutrophils were centrifuged at 350 × g for 5 minutes prior to replacing the supernatant by fresh, serum-free DMEM (for BMDMs) or RPMI (for human neutrophils and macrophages) as control or platelet suspension. For NLRP3 stimulation, cells were primed with 200 ng ml^-1^ LPS for 3 hours and activated with 10 µM Nigericin or 5 mM ATP for 90 minutes unless otherwise indicated. Human monocytes were primed with 2 ng ml^-1^ before being activated with Nigericin (10µM, 90 min). For NLRC4 stimulation, human macrophages were primed with 200 ng/mL LPS for 3 hours before 2 µg ml^-1^ PrgI and 0.5 µg ml^-1^ PA were added to the culture medium for 2 hours. R848 and Pam3cysk4 were also used for priming in some experiments at 10 µM and 1 µg ml^-1^ respectively. After stimulation, cells were centrifuged and supernatants were collected to measure cytokine levels by HTRF. In experiments where the activity of COX1/2 and LOX was inhibited, platelets were incubated with Aspirin, or Zileuton (both at 100 µM) for 60 min at 37°C, before their addition to hMDMs. The drugs were diluted in RPMI 1640, without supplements.

### Real time PCR

Total RNA containing small RNAs from purified human platelets, or PBMCs was purified using the miRNeasy kit (Qiagen), DNA was digested with DNase I (Qiagen), and cDNA was generated by using 500ng RNA with SuperScript III Reverse Transcriptase kit (Thermo Fisher) following the manufacturer’s instructions. cDNA was diluted (1:10for PBMCs, and 1:3 for platelets) and qPCR was performed with Maxima SYBR Green on a Quant Studio 6 Flex RT PCR machine (Thermo Fisher) for 40 cycles and followed by a melt curve analysis for off-target products. The primers used were ACTB-fwd-ccaccatgtaccctggcatt, ACTB-rev-cggagtacttgcgctcagga, NLRP3-fwd-tcggagacaaggggatcaaa, NLRP3-rev-agcagcagtgtgacgtgagg, CD14-fwd-gagctcagaggttcggaaga, CD14-rev-cttcatcgtccagctcacaa, PYCARD-fwd-gagctcaccgctaacgtgct, PYCARD-rev-actgaggaggggcctggat, PF4-fwd-ctgaagaagatggggacctg, PF4-rev-gtggctatcagttgggcagt, CASP1-fwd-acaacccagctatgcccaca, CASP1-rev-gtgcggcttgacttgtccat, GP1BA-fwd-ctgctctttgcctctgtggt, GP1BA-rev-ctccaggtgtgtggtttgtg, IL1B-fwd-tgggcagactcaaattccagct, IL1B-rev-ctgtacctgtcctgcgtgttga.

### Cytokine measurements

Levels of human, or murine IL-1β, IL-6, and TNFα in cell culture supernatants were quantified using commercially available HTRF^®^ (homogeneous time-resolved fluorescence) kits. The HTRF was performed according to manufacturer instructions (CIS bio). Multiplex Cytokine array was used for the detection of human IL-18, IL-1α and MCP1 according to the manufacturer’s instructions (Thermo Fisher).

### Caspase-1 activity

Caspase-Glo^®^ 1 Inflammasome Assay (Promega)provides a luminogenic caspase-1 substrate, Z-WEHD-amino luciferin, in a lytic reagent optimized for caspase-1 activity and luciferase activity.

### Confocal laser scanning microscopy

Platelets and immune cells were imaged in a Leica TCS SP5 SMD confocal system (Leica Microsystems, Wetzlar, Germany). Images were acquired using a 63X objective, with a numerical aperture of 1.2, and analyzed using the Volocity 6.01 software (PerkinElmer, Waltham, Massachusetts, U.S.A.).

### Meta-analysis of microarray data

Pre-processed microarray data of platelets and whole peripheral blood cells under steady state and certain disease states (GSE2006, GSE11524, GSE10361, GSE50858, GSE47018, GSE17924) were down-loaded from the Gene Expression Omnibus (GEO) data base (https://www.ncbi.nlm.nih.gov/geo/). Expression values for the house-keeping gene GAPDH and platelet-as well as inflammasome-associated genes (CXCL4/PFA4, PDGFA, IL1B, PYCARD, NLRP3, TIMP1 CASP1, ARG2, TP2A, SELENB1) were extracted for each data set. In case multiple probe sets for GAPDH were present on the microarray chip, a mean expression value for GAPDH was calculated. The expression values for probe sets of platelet and inflammasome-associated genes were normalized for each data set according the respective GAPDH mean expression value. Log2 transformed normalized expression data were plotted as bar charts by the Tidyverse package in R (v3.4.2).

### Proteomics - Mass spectrometry coupled with liquid chromatography

Suspensions of freshly isolated human platelets were adjusted to a concentration of 5×10^7^/ml in RPMI. Platelets were left untreated, or stimulated with 200 ng/ml LPS, 1 U/ml Thrombin for 3 hours. Platelets were pelleted by centrifugation (3000 × g, 10 minutes) and the cell-free supernatants were harvested. Cell-free supernatants were added with 1x complete protease inhibitor cocktail prior to freezing at −80°C. Proteomics analysis was carried out at the CECAD/CMMC Proteomics Core Facility (University Cologne, Germany) on a Q Exactive Plus Orbitrap (Thermo Scientific) mass spectrometer that was coupled to an EASY-nLC (Thermo Scientific). Briefly, peptides were loaded in 0.1% formic acid in water onto an in-house packed analytical column (50 cm ó 75 µm I.D., filled with 2.7 µm Poroshell EC120 C18, Agilent) and were chromatographically separated at a constant flow rate of 250 nl/minute with the following gradient: 3-4% solvent B (0.1% formic acid in 80 % acetonitrile) within 1 minute, 4-27% solvent B within 119 minute, 27-50% solvent B within 19 minutes, 50-95% solvent B within 1 minutes, followed by washing and equilibration of the columns. The mass spectrometer was operated in data-dependent acquisition mode.

### Data processing and statistical analysis

All mass spectrometric raw data were processed by the CECAD/CMMC Proteomics Core Facility using Maxquant (version 1.5.3.8) with default parameters. Briefely, MS2 spectra were analyzed against the Uniprot HUMAN. fasta (downloaded at: 16.6.2017) database, including a list of common contaminants. False discovery rates on protein and PSM level were estimated by the target-decoy approach to 1% FDR for both. The minimal peptide length was determined to be 7 amino acids and carbamidomethylation at cysteine residues was considered as a fixed modification. Oxidation and acetylation were included as variable modifications. For the analysis, the match-between runs option was enabled. Label-free quantification (LFQ) was activated using default settings. Figures were assembled using the R Tidyverse package.

### Statistical Analysis

Statistical analyses were performed with GraphPad Prism Version 7.0f. Unless indicated otherwise, all graphs are built from pooled data from a minimum of two independent experiments (biological replicates), performed in triplicates (technical replicates). Data are presented as bars and symbols, each symbol representing the average of technical triplicates from individual donors, or mice. The mean and standard deviation (SD) are shown when less than 3, or mean and standard error (SEM) when three or more biological replicates are represented. Additional statistical details are given in the respective figure legends, when appropriate.

### Data availability

A supplementary table containing the source data for all the figure panels will be provided.

---

**Figure S1.**
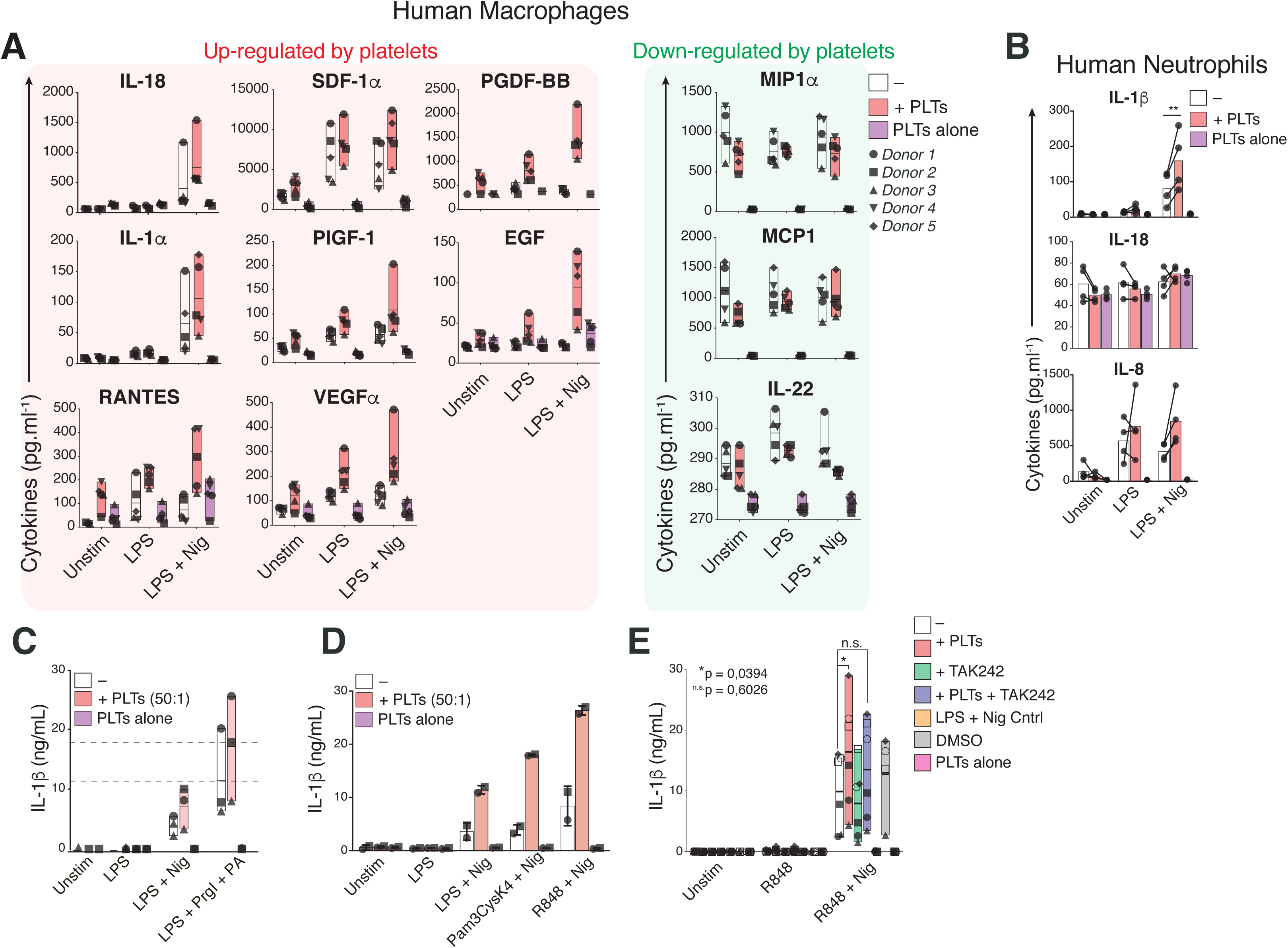
Multiplex measurement of cytokines in cell-free supernatants of unstimulated (Unstim), or LPS-primed (200 ng ml^-1^, 3 hours), and nigericin-activated (10 µM, 90 min) human macrophages (**A**), or human neutrophils (**B**). Cells were cultivated alone (—), or in the presence of platelets (+ PTLs 50:1). (**C**) HTRF measurements of IL-1β in cell-free supernatants of unstimulated (Unstim), or LPS-primed, and PrgI (2 µg ml^-1^) + PA (0.5 µg ml^-1^) stimulated human macrophages that were cultured alone, or in the presence of platelets. (**D**) HTRF measurements of IL-1β in cell-free supernatants of hMDMs primed with LPS, Pam3CysK4 (1 µg ml^-1^), or R848 (10 µM) for 3 hours, followed by activation with nigericin. (**E**) HTRF measurements of IL-1β in cell-free supernatants of hMDMs cultured as in **A**. Cells were pre-treated with TAK242 (0.5 µg ml^-1^) before priming with R848 (10 µM, 3 hours) and activation with nigericin (10 µM, 90 minutes). **A, C** and **E** - Floating bars (with mean and minimum to maximum values) from pooled data from several independent experiments. **B** - Means of one experiment with 4 donors. **D** – Mean ± SD pooled from two independent experiments with platelets and hMDMs from different donors. Each symbol represents the average of technical triplicates of cultures from one donor.

**Figure S2.**
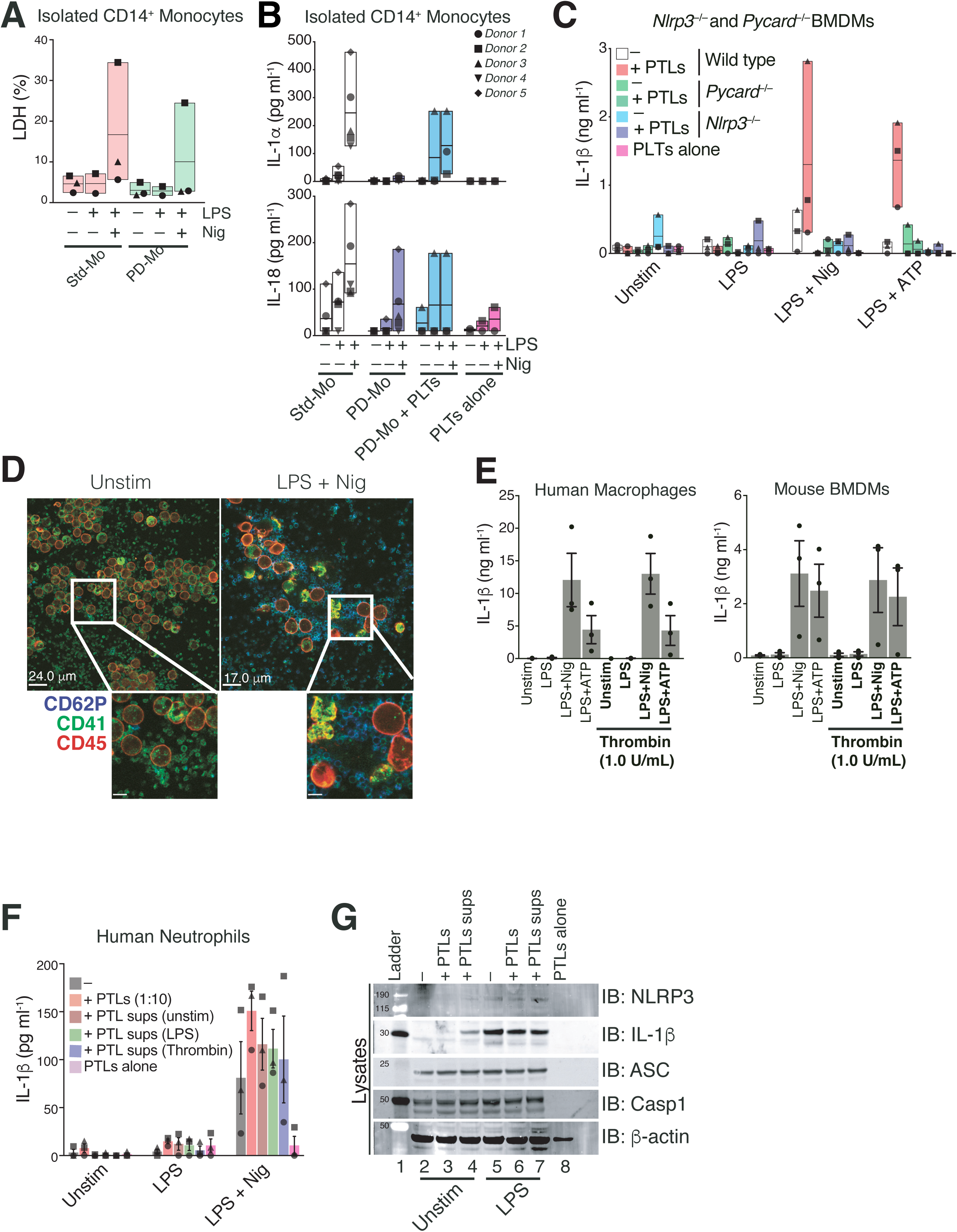
(**A**) LDH release in cell-free supernatants of standard (Std-Mo) or platelet-depleted (PD-Mo) human monocytes. Cells were primed with LPS (2 ng ml^-1^, 3 hours) followed by activation with nigericin (10 μM, 90 min). (**B**) Multiplex Cytokine measurement of IL-1α and IL-18 in cell-free supernatants of Std-Mo, PD-Mo, or PD-Mo that were added with freshly isolated autologous platelets (PD-Mo + PLTs) on a 50:1 platelet-to-monocyte ratio, and platelets cultivated alone. (**C**) IL-1β levels in cell-free supernatants of wild-type, *Nlrp3*^−/−^, or *Pycard*^−/−^ BMDMs cultivated alone (—), or in the presence of wild-type platelets (+ PTLs, 5:1 platelet-to-BMDM ratio). (**D**) Confocal imaging of unstimulated (Unstim), or LPS-primed (200 ng ml^-1^, 3 hours), and nigericin-activated human neutrophils incubated with platelets (50:1 platelet-to-neutrophil ratio). Platelet marker (CD41), platelet activation marker (CD62P), leukocyte marker (CD45). Scale bars are indicated. (**E**) HTRF measurements of IL-1β levels in cell-free supernatants of LPS-primed and nigericin-, or ATP-activated hMDMs (top), or mouse BMDMs (bottom), in the presence or absence of thrombin (1 U ml^-1^). (**F**) IL-1β levels in cell-free supernatants of unstimulated (Unstim), or LPS-primed, and nigericin-activated human neutrophils (10 μM, 90 min) incubated with platelets (10:1 platelet-to-neutrophil ratio) or platelet supernatants from unstimulated, LPS (200 ng ml^-1^, 3 hours) or thrombin (1 U ml^-1^) stimulated platelets. (**G**) Immunoblot of NLRP3, IL-1β, ASC, Caspase-1 and β-actin on whole cell lysates of resting, or LPS-primed human macrophages cultivated alone or in the presence of platelets, or platelet-supernatants. **A - C, F** Floating bars (with mean and minimum to maximum values) from pooled data from 3 - 4 independent experiments. **D** - Representative of two independent experiments. **E** – Mean + SEM from pooled data from three independent experiments. **G** - Representative of 2 independent experiments. Each symbol represents the average of technical triplicates from different donors, or mice.

**Figure S3.**
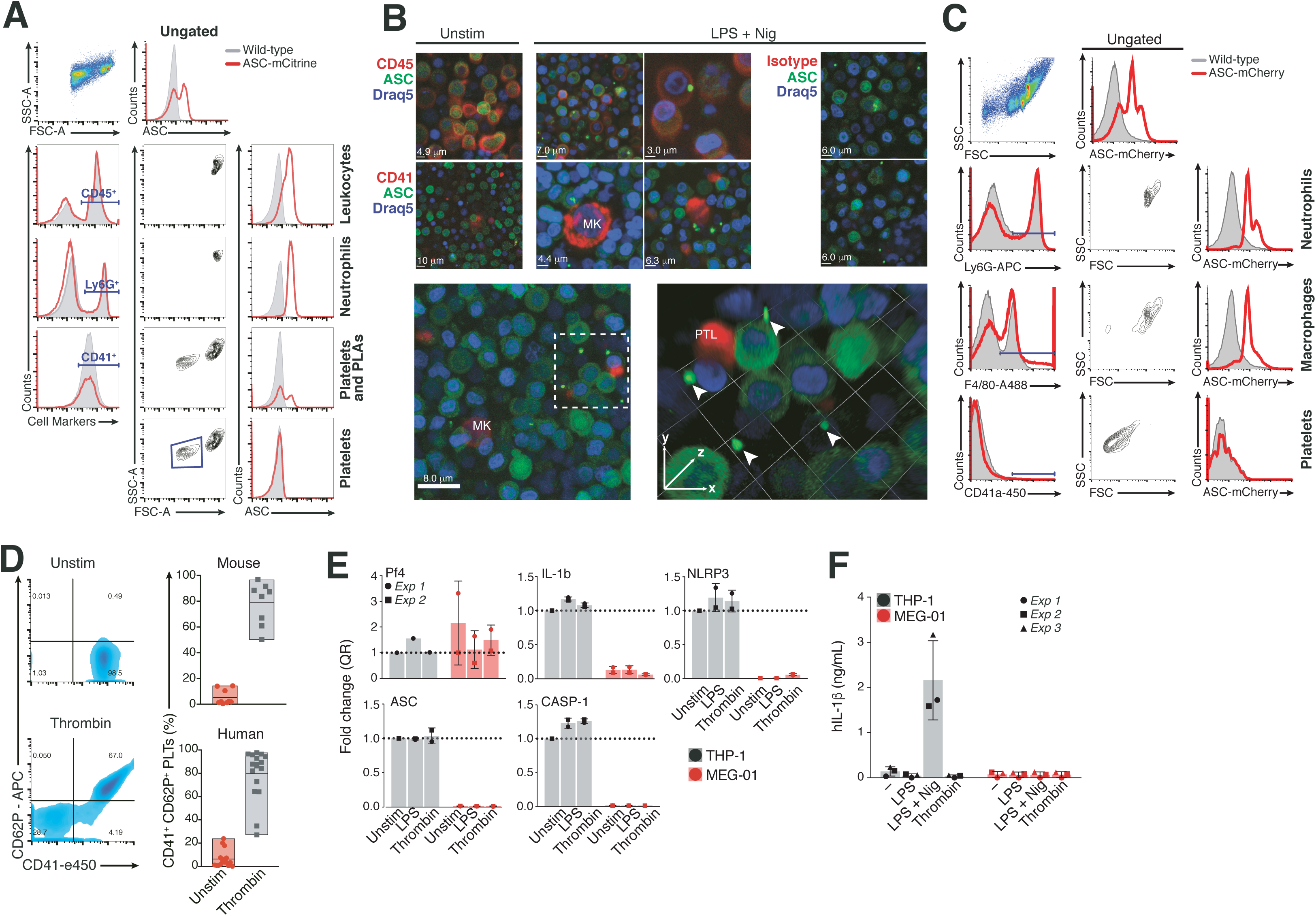
(**A**) Flow cytometric analysis of total bone marrow cells from Wild-type (gray histograms), ASC-mCitrine transgenic, or (**C**) ASC-*knock in* reporter mice (red lines). Gating strategy to identify different cell populations based on surface marker staining and scatter characteristics: CD45 (Leukocytes), F4/80 (Macrophages), Ly6G (neutrophils) and CD41 (platelets and platelet-leukocytes aggregates, PLAs). (**B**) Confocal microscopy of total bone marrow cells from ASC-mCitrine mice, comparing leukocytes (CD45^+^), and platelets (CD41^+^). Megakaryocytes (MKs), platelets (PTL) and ASC specks (arrows). Scale bars are indicated. (**D**) Representative flow cytometric assessment of purity (CD41) and activation (CD62P) of resting, or thrombin (0.5 U ml^-1^) stimulated platelets purified from wild-type mice (n = 8) or human blood donors (n = 18). (**E**) Real time PCR analysis of PF4, IL-1B, NLRP3, ASC and Caspase-1 (CASP-1) expression and (**F**) HTRF measurements of IL-1β in cell-free supernatants on PMA-differentiated THP-1s or the human megakaryocytic cell MEG-01. Cells were left unstimulated, or treated with LPS (1 μg ml^-1^) or Thrombin (1 U ml^-1^). **B** - Representative of two independent experiments. **D** - Floating bars (with mean and minimum to maximum values) from pooled data from several independent experiments, each symbol represents the measurements from platelets of individual mice, or donors. **E** - Mean + SD of pooled data from 2 independent experiments performed in triplicates.

**Extended Data Fig. 4.**
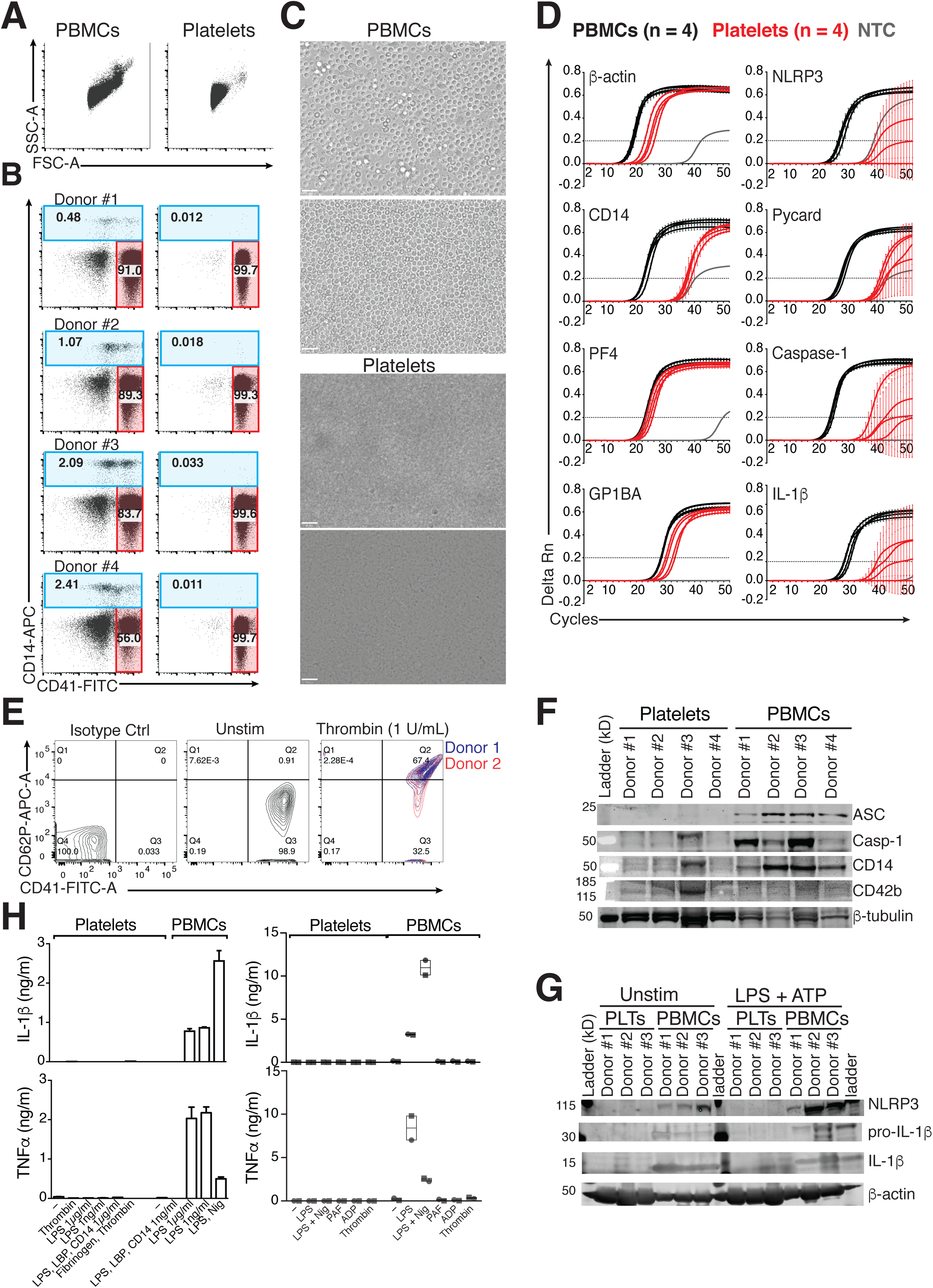
(**A-B**) Representative flow cytometric analysis of PBMCs, or platelets purified from peripheral blood of healthy donors (n = 4) showing the purity of platelet preparations with CD14 (monocyte) or CD41 (platelets) surface markers. (**C**) Light microscopy of PBMCs, or isolated platelets. Scale bars: 100 μm. (**D**) Amplification plots of the expression of NLRP3, PYCARD, Caspase-1, and IL-1B, in comparison to Monocyte marker (CD14) and platelet markers (PF4 and GP1BA) from a qPCR performed on total RNA isolated from PBMCs or platelets (n = 4). (**E**) Flow cytometric assessment of purity (CD41) and activation (CD62P) of resting, or thrombin (1 U ml^-1^) stimulated platelets purified from healthy donors (n = 2). (**F**) Immunoblotting of ASC, Caspase-1, CD14, CD42 and β-tubulin in resting platelets, or PBMCs from healthy donors (n = 4). (**G**) Immunoblotting of NLRP3, or β-actin in resting, or activated platelets, or PBMCs from healthy donors (n = 3). (**H**) IL-1b levels in cell-free supernatants of human isolated Platelets or PBMCs that were stimulated as indicated for 3h (left), or 18h (right) (n = 2). **A - D, F** - Representative of two independent experiments performed with 4 healthy blood donors. **E** - Representative of several experiments. Two donors are shown. **G** - Data from one experiment with 3 different donors. **H** – Mean + SD of technical triplicates. Representative of 2 independent experiments.

**Figure S5.**
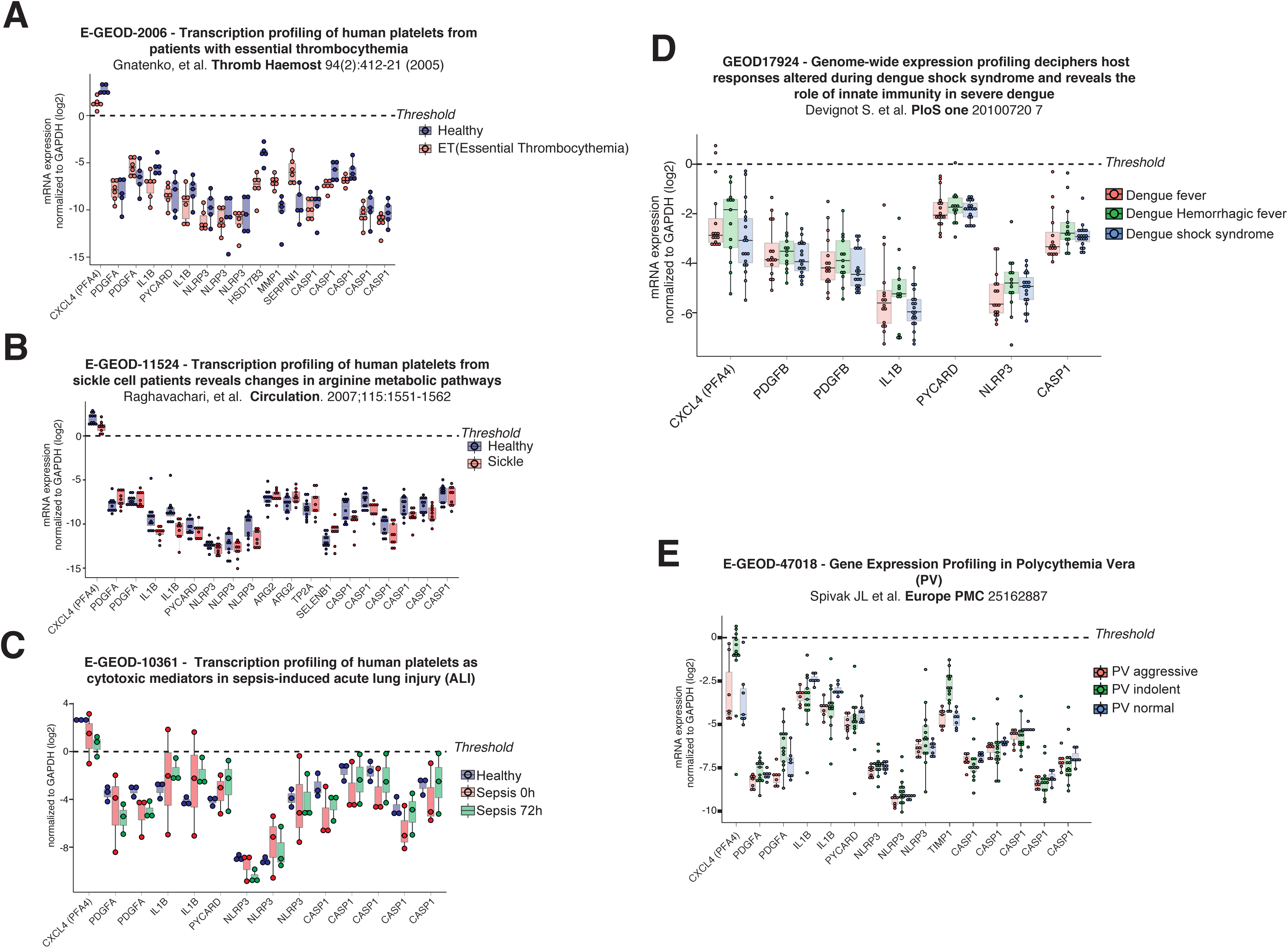
**A – E** Microarray Meta-analysis of transcriptomics of isolated human platelets. Gene expression data from five independent studies that addressed the transcriptomics of platelets isolated from the peripheral blood of healthy donors, or patients with a variety of diseases. Bar charts represent the Log_2_ transformed data for platelet marker (CXCL4, and PDGFA) as well as inflammasome-associated genes (PYCARD, NLRP3, TIMP1 CASP1, ARG2, TP2A, SELENB1). Expression values were normalized against the expression of the constitutive gene GAPDH.

**Figure S6.**
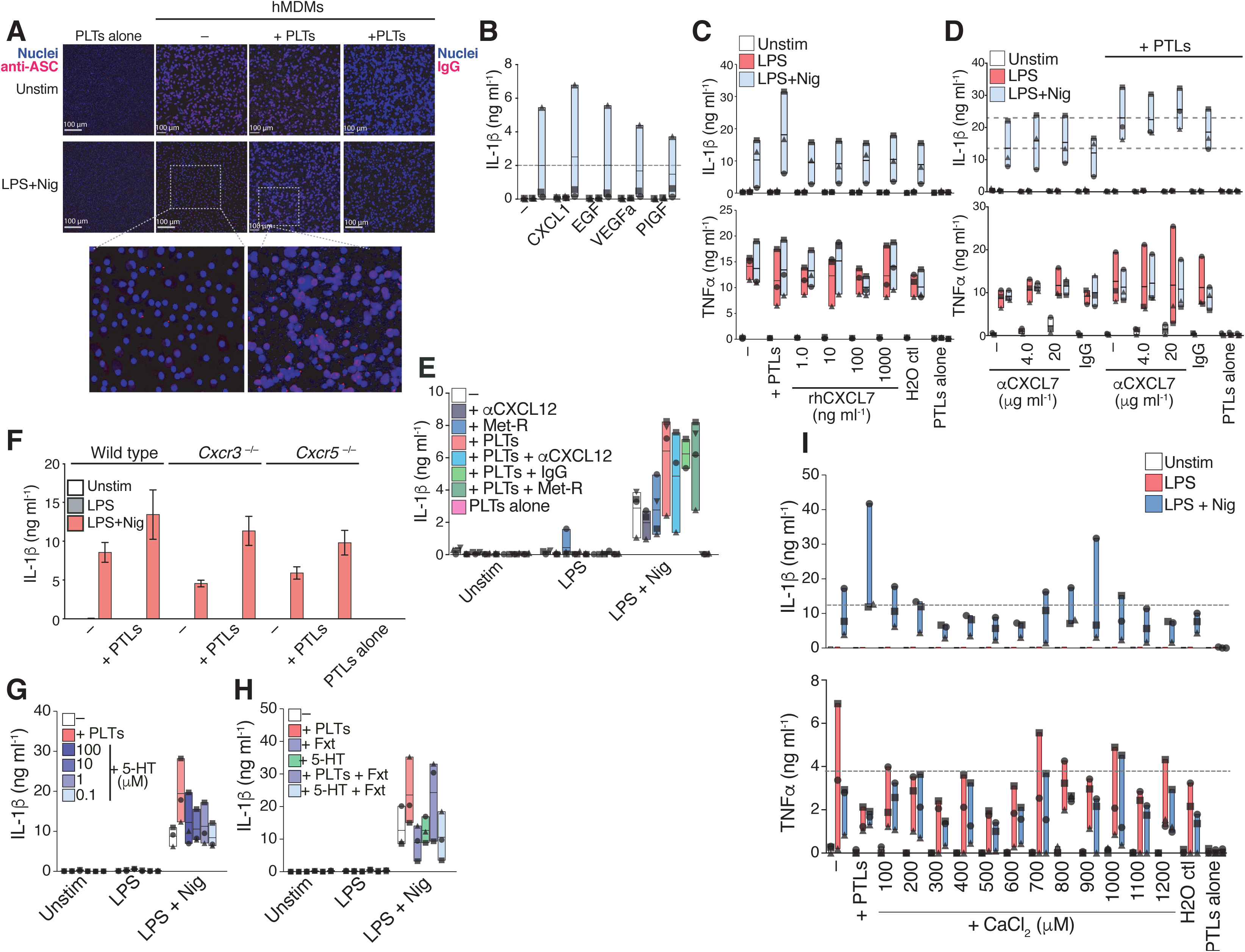
(**A**) Confocal microscopy of ASC specks in LPS primed (200 ng ml^-1^, 3 hours) and nigericin activated (10 μM, 45 min) human macrophages. Cells were either cultured alone (−) or in the presence of platelets (+PLTs, 50:1 platelet-to-macrophage ratio). (**B - e**) IL-1β and TNFα levels in cell-free supernatants of unstimulated (−), LPS stimulated, and nigericin-activated (10 μM, 90 min) human macrophages, either cultivated alone (−), or in the presence of platelets (+ PTLs, 50:1 platelet-to-macrophage ratio) as indicated. (**B**) Addition of the following recombinant human proteins to human macrophages: CXCL1 (100 pg ml^-1^), EGF (200 pg ml^-1^), VEGFa (200 pg ml^-1^), PIGF (80 pg ml^-1^) or (**C**) the indicated concentrations of CXCL7. (**D**) Blocking antibodies against CXCL7 (4 or 20 µg ml^-1^) or (**E**) against CXCL12 (10 µg ml^-1^) added to macrophages before the start of the assay. To block CCR5, human macrophages were pre-incubated with met-RANTES (10 ng ml^-1^, 90 minutes) before the start of the assay. Matching IgG isotype controls were added at the same concentrations as the respective blocking antibodies. (**F**) IL-1b levels in cell-free supernatants of unstimulated (−), LPS stimulated, and nigericin-activated wild-type, *Cxcr3*^−/−^, or *Ccr5*^−/−^ BMDMs cultivated alone (−), or in the presence of wild-type platelets (+ PTLs, 5:1 platelet-to-BMDM ratio). (**G** - **H**) IL-1b levels in cell-free supernatants of hMDMs stimulated as in B, and added with recombinant human serotonin (5-HT) at the indicated concentrations (**G**), or pre-incubated with fluoxetin (10 mm) for 30 min before the addition of platelets. (**I**) IL-1b and TNFa levels in cell-free supernatants of unstimulated (−), LPS stimulated, and nigericin-activated hMDMs cultivated alone (−), or in the presence of platelets (+ PTLs, 50:1 platelet-to-macrophage ratio), added with the indicated concentrations of calcium chloride (CaCl_2_) in Ca^2+^-free medium before the start of the assay. **A** - Representative of three independent experiments **B** – **E, G - I** Floating bars (with mean and minimum to maximum values) from pooled data from independent experiments. Each symbol represents the average of technical triplicates from different donors. Representative of three - four independent experiments. **F** - Mean + SD of technical triplicates from one experiment.

**Figure S7.**
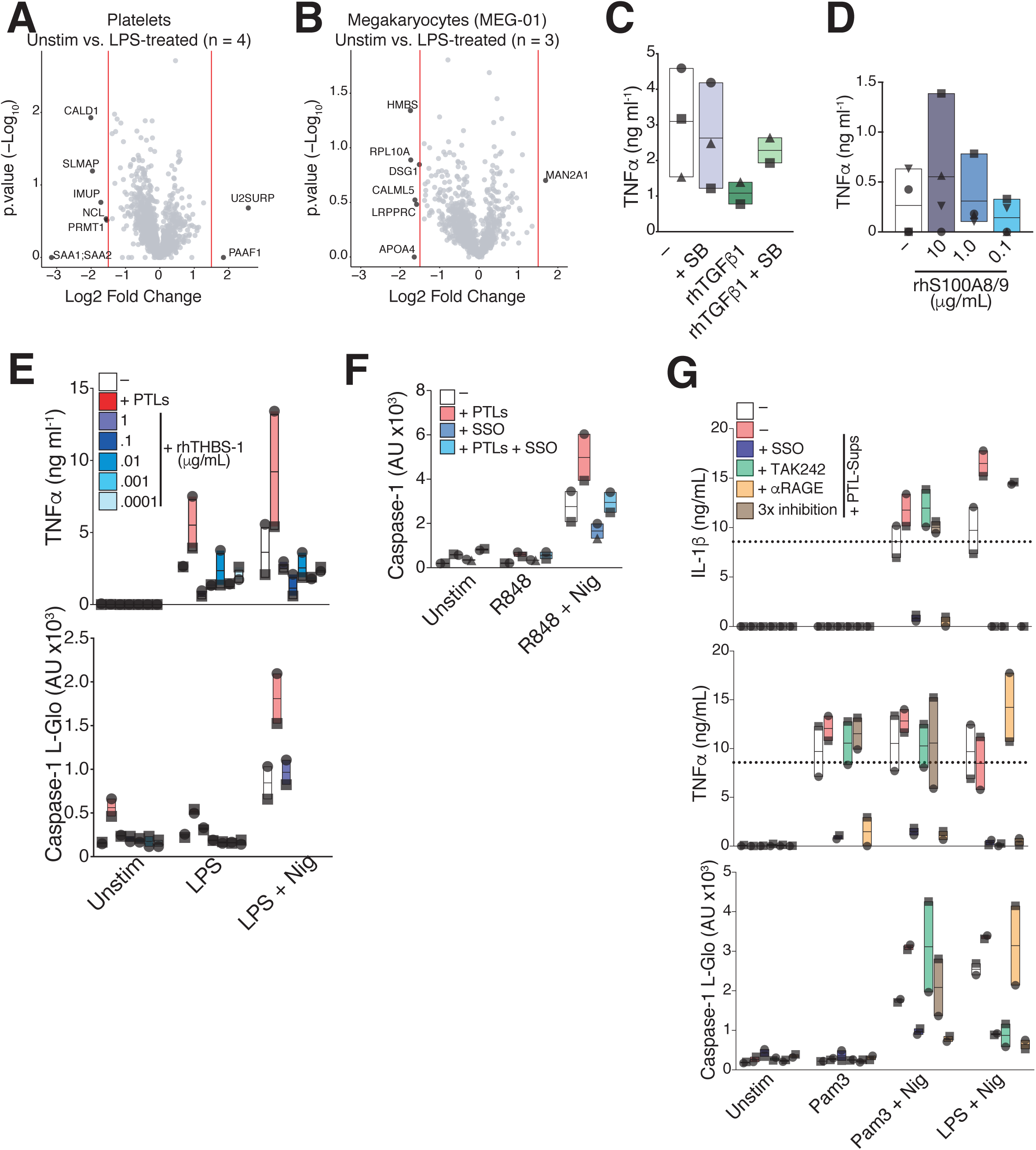
(**A - B**) Volcano plots showing the Log2-fold change and −Log10 of the p values of secretome of (**A**) human platelets (n = 4) and (**B**) megakaryocytes (n = 3). Proteins with Log2 fold >= 2 | <= −2 are labeled. (**C**) TNFα levels in cell-free supernatants of hMDMs that were left untreated or pre-treated with the TGFb-inhibitor SB-431542 (10 μM, for 1 hours), followed by stimulation with recombinant human TGF-b1. (**D**) TNFα levels in cell-free supernatants of hMDMs that were left untreated or stimulated with recombinant human S100A8/9 at the indicated concentrations. (**E**) TNFα levels and caspase-1 activity in cell-free supernatants of unstimulated, or LPS-primed hMDMs that were co-incubated with platelets, or the indicated concentrations of rhTHBS1. (**F**) Caspase-1 activity measured in cell-free supernatants of unstimulated, or R848-primed hMDMs pre-treated or not with the CD36 inhibitor SSO. (**G**) IL-1b, TNFa levels and caspase-1 activity in cell-free supernatants of unstimulated, or LPS-or Pam3Cys-K4-primed hMDMs that were activated with nigericin in the presence of SSO (50 µM), TAK242 (0.5 µg ml^-1^), anti-Rage (10 µg ml^-1^), or all those inhibiting strategies combined (3X inhibition). **C - G** - Floating bars (with mean and minimum to maximum values) from pooled data from a minimum of 2 independent experiments. Each symbol represents the average of technical replicates from different donors.

